# Age-dependent dormant resident progenitors are stimulated by injury to regenerate Purkinje neurons

**DOI:** 10.1101/263954

**Authors:** N. Sumru Bayin, Alexandre Wojcinski, Aurelien Mourton, Hiromitsu Saito, Noboru Suzuki, Alexandra L. Joyner

## Abstract

Outside of the neurogenic niches of the brain, postmitotic neurons have not been found to undergo efficient regeneration. Here we demonstrate that Purkinje cells (PCs), which are born at midgestation and are crucial for both development and function of cerebellar circuits, are rapidly and fully regenerated following their ablation at birth. New PCs are produced by a previously unidentified progenitor population and support normal cerebellum development. The number of PC progenitors and their regenerative capacity, however, diminish soon after birth, and consequently PCs are poorly replenished when ablated at postnatal day 5. Nevertheless, the PC-depleted cerebella reach a normal size by increasing cell size, but scaling of neuron types is disrupted and cerebellar function is impaired. Our findings thus provide a new paradigm in the field of neuron regeneration by identifying a unipotent neural progenitor that buffers against perinatal brain injury in a stage-dependent process.

**One sentence summary:** Injury induces a dormant progenitor population present at birth to regenerate cerebellar neurons in a time-dependent manner.

## Introduction

Most neurons in the brain are generated at specific developmental time points, and once a neuron is postmitotic regeneration following injury is limited, except for in two forebrain regions that maintain neurogenesis (Chaker, Codega, & Doetsch, 2016). In the context of injury, adult forebrain neurons undergo limited recovery that involves either reactive gliosis (Buffo et al., 2008; Robel, Berninger, & Gotz, 2011; Sirko et al., 2013) or migration of neural stem cells from the neurogenic niches (Benner et al., 2013; Llorens-Bobadilla et al., 2015; Lopez-Juarez et al., 2013; Marti-Fabregas et al., 2010). The cerebellum (CB) of the hindbrain has a complex folded structure that houses the majority of neurons in the brain and is essential for balance and motor coordination, as well as higher order reasoning via circuits it forms throughout the forebrain (Fatemi et al., 2012; Steinlin, 2007; Tavano et al., 2007; Tsai et al., 2012; Wagner, Kim, Savall, Schnitzer, & Luo, 2017). The postnatal developing mouse CB maintains two neurogenic progenitor pools for two weeks. Interestingly, the proliferating granule cell progenitors were recently found to be replenished following injury by adaptive reprograming of the second Nestin-expressing glial progenitors (Wojcinski et al., 2017). However, once a neurogenic process has ended, the degree to which post mitotic neurons can undergo regeneration is poorly understood.

Purkinje cells (PC) are born by embryonic day (E) 13.5 in the mouse and during weeks 10-11 in humans (Rakic & Sidman, 1970; V. Y. Wang & Zoghbi, 2001). After exiting the cell cycle in the ventricular zone, PCs express FOXP2 and migrate to form a PC layer (PCL) under the cerebellar surface by E17.5, and turn on Calbindin1 (CALB1) as they mature. PCs play a central role in CB development by being the main source of sonic hedgehog (SHH), which is required for proliferation of granule cell progenitors and Nestin-expressing progenitors (Corrales, Blaess, Mahoney, & Joyner, 2006; Fleming et al., 2013; Lewis, Gritli-Linde, Smeyne, Kottmann, & McMahon, 2004). PCs also are key for CB function by integrating the inputs that converge on the cerebellar cortex (Sillitoe & Joyner, 2007). Hence, PC loss is linked to cerebellar motor behavior syndromes and has also been implicated in autism (Fatemi et al., 2012; Tsai et al., 2012; S. S. Wang, Kloth, & Badura, 2014). It is thus essential to determine the regenerative potential of PCs around birth.

## Results and Discussion

To ablate and track PCs, the diphtheria toxin receptor (DTR) and a lineage tracer, TdTomato (TdT), were expressed in PCs (*Pcp2^Cre/+^; R26^LSL-DTR/LSL-TdT^* or *PC-DTR* mice; LSL = lox-stop-lox). At postnatal day (P) 1, 53.1 ± 22.6% of PCs (n = 5 mice) expressed TdT, and all TdT+ cells expressed CALB1 and DTR (Figure 1_Supplement 1). Strikingly, when DT was injected at P1 into *PC-DTR* pups (P1-*PC-DTR)*, nearly all TdT+ PCs formed an ectopic inner layer by 1 day post injection (dpi) (Figure 1A-M). The ectopic layer was absent by P8 (Figure 1 K), and TdT+ cells in the ectopic layer were TUNEL positive starting at P3 with a peak at P5, indicating that almost all DTR-expressing TdT+ cells became misplaced, died and were cleared within 5-7 dpi of DT (Figure 1N,O).

**Figure 1.**
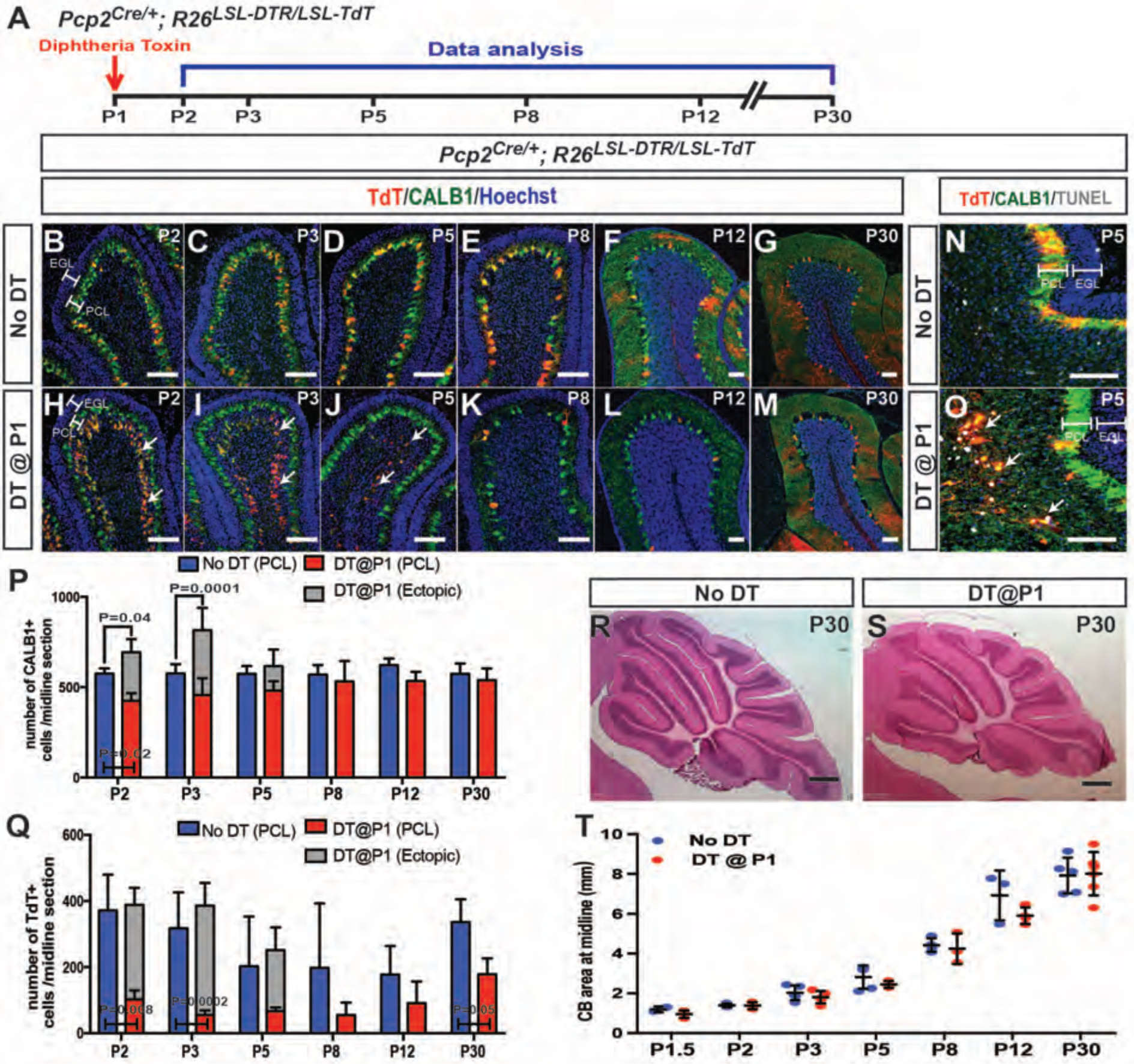
P1 DT-ablated PCs are replenished and CB size and morphology appears normal. A. The experimental plan. **B-M.** IF analysis at the indicated ages for TdT and CALB1 in sagittal cerebellar sections of lobule IV-V in No DT (b-g) and P1-*PC-DTR* mice (h-m). **N-O**. Analysis of apoptosis at P5 using TUNEL. **P**. Quantification of CALB1+ cells per midline section in PCL (blue or red) and ectopic layer (grey) (PCL cells: Two-way ANOVA F_(5,49)_=3.586, P = 0.008, and total number of PCs: Two-way ANOVA F_(5,27)_=4.732, P = 0.003, n ≥ 3 animals/condition) **Q.** Quantification of TdT+ cells per section (PCL cells: Two-way ANOVA F_(5,43)_=7.22, P = 0.0001). Significant *post hoc* comparisons are shown. **R-S.** H&E stained midline sagittal sections of cerebella at P30 of No DT (R) and P1-*PC-DTR* (S) mice. **T**. Quantification of midline sagittal areas of cerebella shows no differences upon DT injection (P = 0.89, n ≥ 3 for each age) Scale bars: 200 µm, (R-S) 500 µm. (EGL: external granule layer, PCL: Purkinje cell layer)

Unexpectedly, although the number of CALB1+ PCs in the PCL of P1-*PC-DTR* mice was significantly reduced at P2 compared to non-injected controls (No DT), it was not reduced at P3 and later stages (Figure 1P). Furthermore, the total number of PCs (ectopic layer + PCL) was significantly greater in DT-injected cerebella than in No DT controls at P2 and P3, and the total number of PCs was down to normal levels at P5, overlapping with the time of clearance of the ectopic layer (Figure 1P). The number of TdT+ cells in the PCL remained significantly lower in P1-*PC-DTR* brains at P30 compared to No DT controls (Figure 1Q). Consistent with the rapid recovery of PC numbers in the PCL, no significant decrease in the area of the CB on sections was observed at between P1.5 and P30 (Figure 1R-T Figure1_supplement 2), or in the thickness of the outer (proliferating) and inner (differentiating) external granule cell layers (Figure1_supplement 3). In summary, we uncovered that the CB can rapidly recover (within 24 h) from the loss of ∼50% of PCs at P1, by producing new PCs and resuming normal growth.

In order to document the rapid production of new PCs after ablation, we tested whether PCs that had recently undergone cell division could be detected at P3. P1-*PC-DTR* mice were divided into 4 groups; each group receiving three injections of BrdU (2 h apart) during 4-26 h after DT-injection (Figure 2A). Indeed, BrdU+ cells were observed in the PCL of all groups (Figure 2B), and apart from astrocytes and microglia (Figure2_supplement 1), all were PCs (FoxP2+ and CALB1+), with the greatest incorporation being between 10-20 h after DT (Figure 2B,E,F). Importantly, no BrdU incorporation was observed in PCs in No DT mice (Figure 2C,D). Furthermore, a lack of BrdU incorporation in the ectopic layer confirms that the labeling is not due to DNA damage induced by DT-mediated cell death (Figure 2E,F). These results reveal that a progenitor capable of proliferating produces the new PCs after ablation at P1.

**Figure 2.**
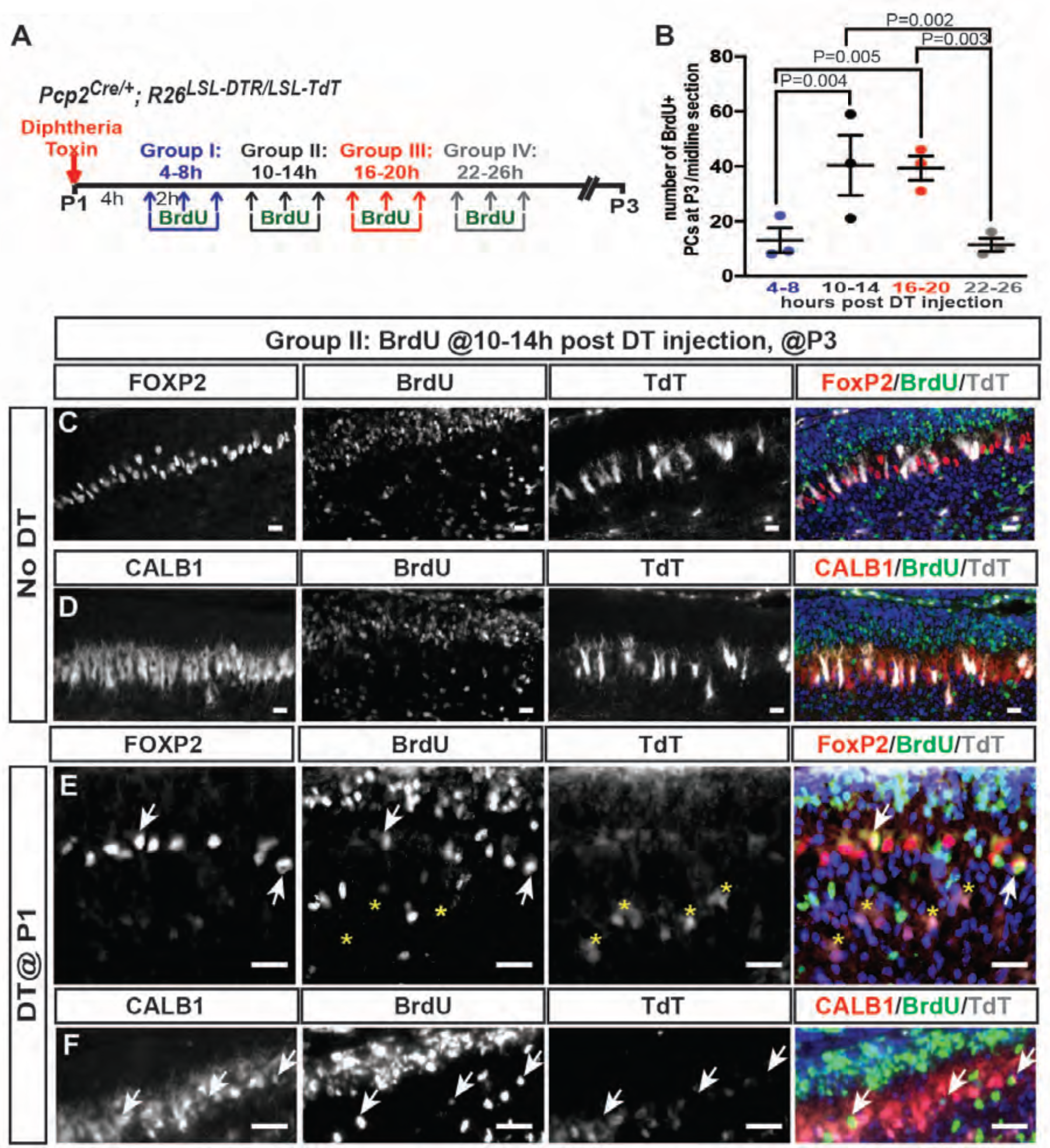
Progenitors proliferate within 24 hours of DT-injection at P1 in *PC-DTR* mice and produce new PCs. A. The experimental plan. **B.** Quantification of the number of BrdU+ PCs (CALB1+) at P3 in P1-*PC-DTR* mice (Two-way ANOVA F_(3,16)_=6.163, P = 0.006, n = 3 animals/condition). Significant *post hoc* comparisons are designated in the figure. **C-F.** Representative images from BrdU injection performed 10-14 h post DT injection at No DT (C, D) or P1-*PC-DTR* (E, F) or mice P3 cerebella were shown (n = 3 animals/condition). IF analysis at P3 from No DT brains showed no BrdU incorporation in PCs, identified by either FOXP2(C) or CALB1(D). IF analysis of P1-*PC-DTR* animals showing FoxP2+ (E) and CALB1+ (F) and BrdU+ cells (arrows) at P3. Asterix shows TdT+ cells are BrdU-. Scale bars: 50 µm

Based on the rapid response to ablation, we hypothesized that local progenitors in the PCL are responsible for the PC replenishment. The population of Nestin-expressing progenitors in the PCL was a candidate, as they display plasticity upon ablation of granule cell precursors in newborn mice (Wojcinski et al., 2017). Furthermore a putative rare Nestin+ cell in the adult CB was recently described to produce new neurons in response to exercise (Ahlfeld et al., 2017). However, when we tested the contribution of Nestin-expressing progenitors to PC regeneration using a *Nes-CFP* reporter allele that transiently maintains CFP protein after differentiation, no CFP+ cells were found to co-express FOXP2 or CALB1 at 12 h and 2 days post DT injection in P1-*PC-DTR* mice and in No DT controls (Figure2_supplement 2). These results suggested that another progenitor population mediates regeneration following PC depletion.

We next examined whether a progenitor exist after birth that express early (FOXP2) but not late (CALB1) PC markers. Indeed, at P1 we detected cells in the PCL that expressed FOXP2 but showed only low or no CALB1 expression (named FEPs for FOXP2-expressing progenitors; Figure 3A-B). Furthermore, temporal analyses revealed a steady decrease in the number of FEPs from P1 (74.33 ± 5.69/midline sagittal section) to P5 (28.66 ± 7.51/midline sagittal section, Figure 3A,C), revealing the progenitors are a transient population. Interestingly, the few FEPs present at P5 were specifically enriched in the central and nodular zones of the CB, which are developmentally delayed at P5 (Legue, Riedel, & Joyner, 2015; Sudarov & Joyner, 2007)(Figure 3A). In addition, FEP numbers significantly increased 12 hours after DT injection in P1-*PC-DTR* mice (1.90 ± 0.05-fold, Figure 3D), indicating they expand upon injury as the expansion correlates with the highest BrdU incorporation time window we observed after injury (Figure 2B). In addition, using *FoxP2^Flpo/+^; R26^FSF-TdT/+^* (FSF = frt-stop-frt) mice in which all PCs and FEPs express TdT at P1 (Figure3_supplement 1), an increase in transiently fate mapped TdT*+* FEPs was observed 12 hours after DT injection at P1 (1.86 ± 0.46–fold, n = 3, Figure3_supplement 2). Surprisingly, at P5 the number of FEPs was significantly lower in P1-*PC-DTR* animals than in No DT mice (Figure 3D), possibly reflecting an exhaustion of the progenitor population by production of new PCs.

**Figure 3.**
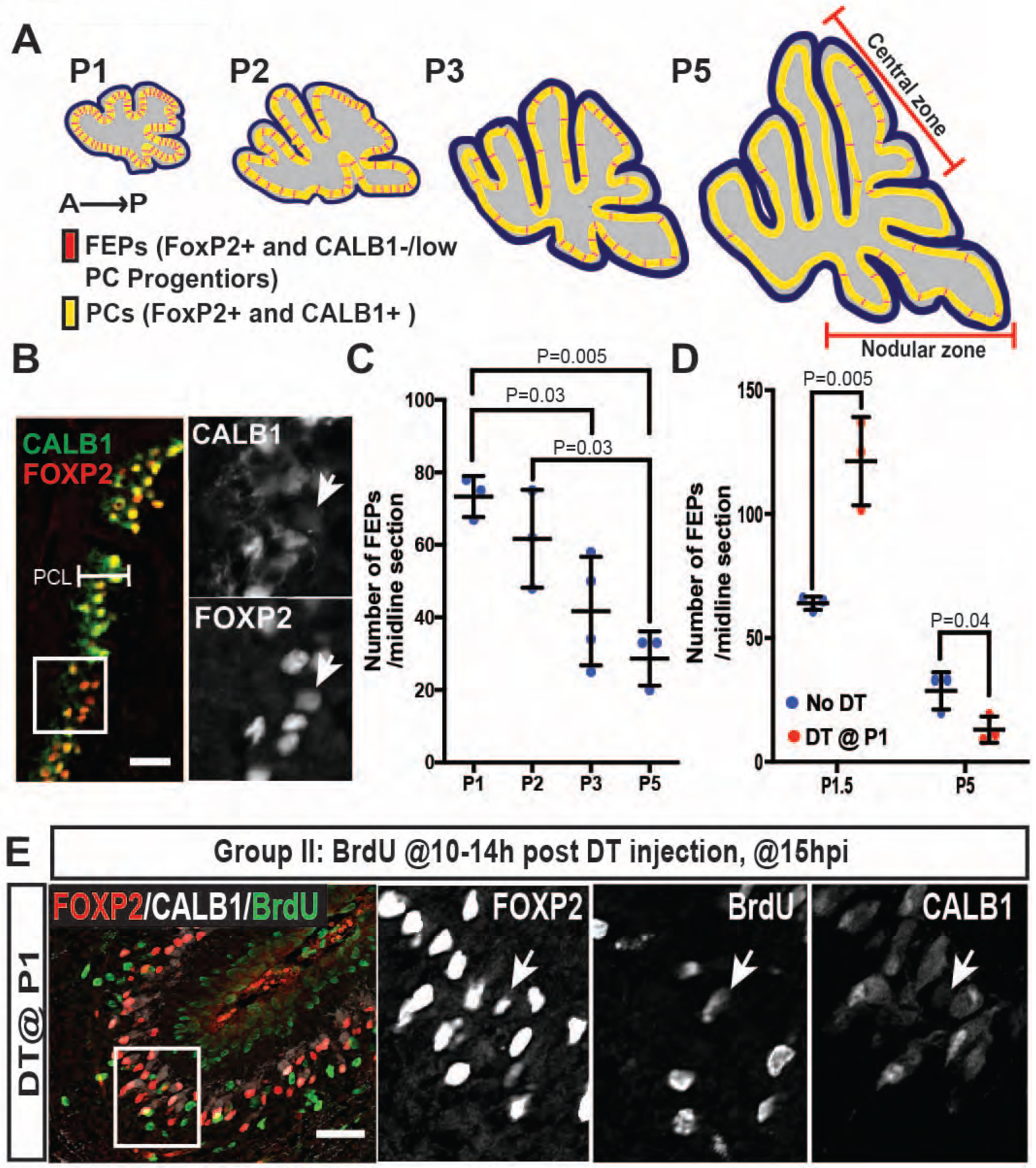
Number of FEPs diminishes with age and increases after ablation of PCs. **A.** Schematic representation of the distribution of FEPs (red) in sagittal midline sections of P1-5 cerebella (yellow, FoxP2+ and CALB1+ PCs) **B.** IF analysis of FEPs (FoxP2+ and CALB1-/low, arrow) at P1.5 in No DT-injected mice. **C.** Quantification of the numbers of FEPs at P1-5 (One-way ANOVA F_(3,9)_=9.074, P = 0.004, n ≥ 3 animals/condition). Significant *post hoc* comparisons are shown. **D.** Quantification of the numbers of FEPs at P1.5 (Two-tailed t-test, P = 0.005, n = 3) and P5 (Two-tailed t-test, P = 0.04, n = 3) in No DT and P1-*PC-DTR* mice. **E.** Arrow shows a BrdU+ FEP (CALB1-/low, FoxP2+) at 15 h post injection (hpi) in P1-*PC-DTR* mice (n = 3). Scale bars: 100 µm

To confirm that FEPs undergo proliferation upon PC depletion, we injected BrdU 10-14 h after DT and collected cerebella 1 h (∼P1.5) later. All BrdU+ PCs in the PCL of P1-*PC-DTR* mice expressed FOXP2, but only 45.5 ± 1.1% expressed CALB1 (Figure 3E, Figure3_supplement 3). In addition, Ki67+ FoxP2+ cells were detected at P1.5 in the PCL of P1-*PC-DTR* pups (Figure3_supplement 3C), confirming the presence of proliferative FEPs following PC ablation. Furthermore, the total number of BrdU+ PCs in the PCL was the same at P1.5 (38.7 ± 9.1/section, n = 3) and at P3 (40.3 ± 19.0/section, n = 3). Collectively, our data argues that the recovery of PCs in P1-*PC-DTR* mice is mediated by a previously unrecognized dormant (Ki67- and not labeled with BrdU) and age-dependent progenitor population (FEPs) that proliferates and differentiates into PCs in response to injury.

Given that the population of FEPs is greatly reduced by P5 (Figure 3C), PCs should not be efficiently replaced when ablated at P5. Indeed, when DT was injected at P5 (P5-*PC-DTR* mice) (Figure 4_supplement 1A), the numbers of PCs were significantly reduced by P12 compared to No DT controls (Figure 4_supplement 1B-I, R). TdT+ PCs were TUNEL+ by P8 (Figure 4_supplement 1J-K) and the majority of TdT+ cells were cleared from the PCL by P12 (Figure 4_supplement 1G, P and S). Furthermore, PCs had abnormal dendrites at P8 and P12 (Figure 4_supplement 1B-G and L-P) and some PCs had misplaced somas at P30 (Figure 4_supplement 1N-R). Interestingly, the reduction in PCs numbers observed by P12 was maintained at P30 (Figure 4_supplement 1R), such that the number of PCs was reduced by 32.4 ± 6.5%. In summary, there is little replenishment of PCs when they are ablated at P5 (Figure 4A).

**Figure 4.**
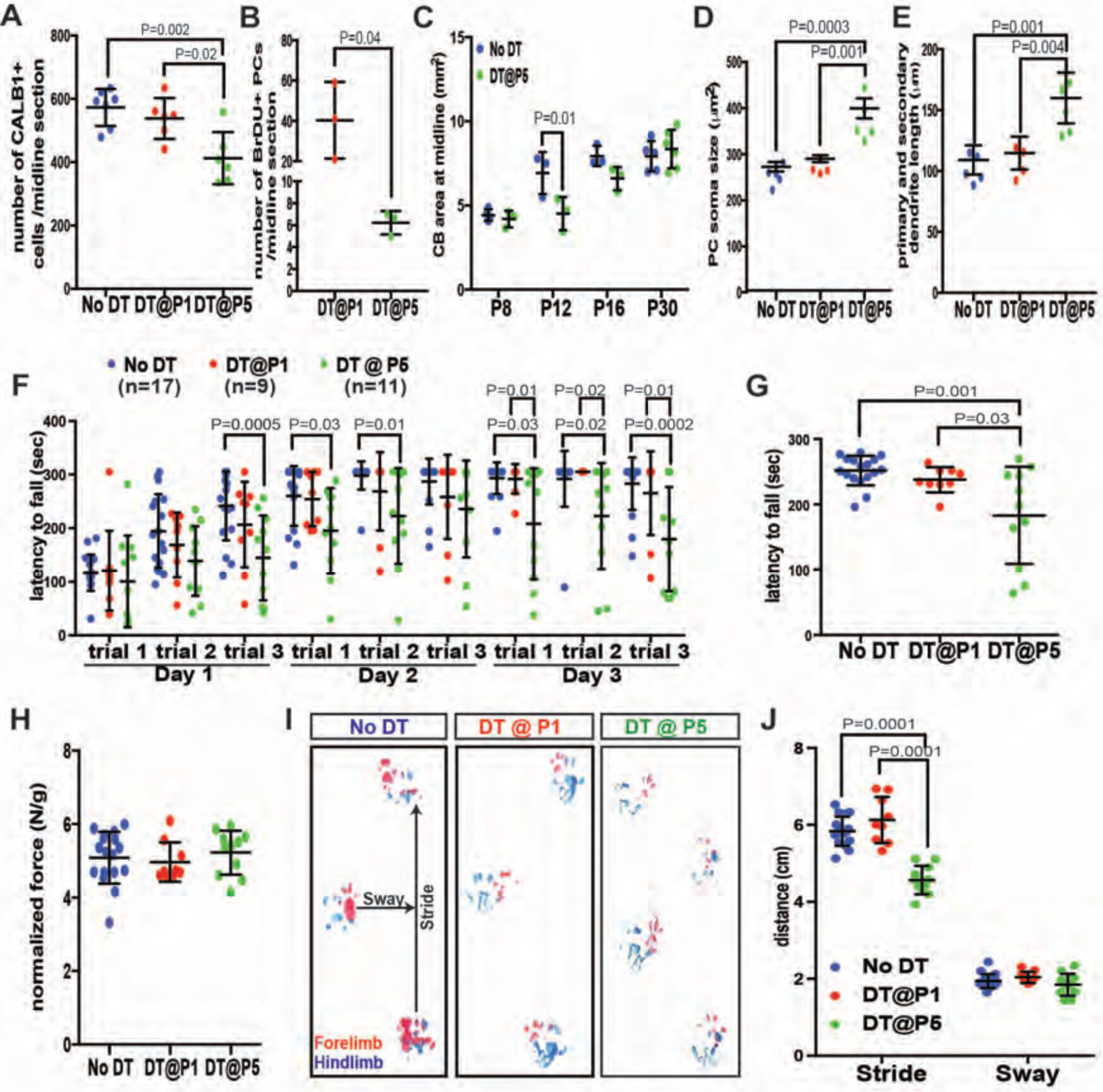
Despite the recovery of CB size, PC are poorly replenished and altered and motor behavior deficits develop when PCs are killed P5 but not at P1. **A.** Number of CALB1+ cells at P30 (One-way ANOVA, F_(2,16)_=9.464, P = 0.002, n ≥ 6). **B.** Number of BrdU+ PCs 2 days post DT-injection at P1- or P5- *PC-DTR* mice (Two-tailed t-test, P = 0.04). **C**. Quantification of CB area in midline sagittal sections demonstrates that CB size is smaller at P12 in P5-*PC-DTR* mice but not later (Two-way ANOVA, F_(1,22)_=7.045, P = 0.01, n ≥ 3). **D-E.** PC soma size (D, One-way ANOVA, F_(2.11)_=20.56, P = 0.0002, n ≥ 4) and primary and secondary dendrite lengths (E, One-way ANOVA, F_(2,11)_=14.54, P = 0.0008, n ≥ 4) at P30 were increased in P5*-PC-DTR* animals compared to No DT and P1-*PC-DTR* animals. **F-G.** Latency to fall from rotarod at each trial (F, Two-way ANOVA, F_(2,34)_=8.37, P = 0.001, n ≥ 9) and cumulative analysis (G, One-way ANOVA, F_(2,34)_=11.12, P = 0.0002, n ≥ 9, No DT vs. DT@P1: P = 0.83). **H.** Analysis of grip strength showed no change in P1 (n = 9, vs No DT: P = 0.89) and P5 (n = 11, vs. No DT: P = 0.84, vs. DT@P1: P = 0.64) DT-injected mice compared to controls (No DT, n = 17). **I-J.** Representative images (I) and quantification (J) of footprint analysis performed on P1- (vs. No DT: stride: P = 0.10 and sway: P = 0.90) and P5-*PC-DTR* mice and controls (Two-way ANOVA, F_(2,133)_=73.45, P = 0.0001, n ≥ 9). Significant *post hoc* comparisons are shown.

We next tested whether the rare FEPs at P5 (Figure 3A,C) can still proliferate upon PC depletion. Unlike in P1-*PC-DTR* mice, very few BrdU+ FEPs were detected in P5-*PC-DT*R cerebella injected with BrdU at 10-14 h post DT-injection at both 1 h (5.55 ± 0.51/ midline sagittal section, n = 3) and 1.5 days (6.22 ± 1.07/ midline sagittal section, n = 3, Figure 4B) post BrdU-injection and the few BrdU+ FEPs were concentrated in the central and the nodular zones enriched for FEPs at P5 (Figure 4_supplement 2). Interestingly, compared to P1-*PC-DT*R mice in which 52.29 ± 0.09% (n = 3) of FEPs incorporated BrdU, only 20.55 ± 0.07% (n = 3) incorporated BrdU in P5-*PC-DTR* animals. Overall, these results demonstrate that replenishment of PCs is not efficient at P5 because with age, FEPs both diminish in number and in their ability to proliferate in response to PC depletion.

We next examined whether the depletion of PCs in P5-*PC-DTR* mice had an effect on CB development. Indeed, the area of CB sections was significantly reduced at P12 but not P8 (Figure 4C), and the thickness of the external granule cell layer was significantly reduced in P5-*PC-DTR* mice at P8, but by P12 the decrease was diminished (Figure4_supplement 4A-E). However, despite the lack of recovery of PC numbers the reduction in CB size became less pronounced with age and by P30 the area of the CB was normal (Figure 4C, Figure4_supplement 3A-I). As a consequence there was a reduction in PC density compared to No DT or to P1-*PC-DTR* mice (Figure4_supplement 3J, Figure4_supplement 2N,Q). The density of granule cells also was lower compared to No DT and P1-*PC-DTR* P30 mice (Figure4_supplement 4F). Interestingly, PCs in P5-*PC-DTR* mice had a larger soma (Figure 4D) and longer primary and secondary dendrites (Figure 4E) compared to No DT or P1-*PC-DTR* mice, a cellular phenotype observed in some mouse mutants with PC loss (Castagna, Merighi, & Lossi, 2016). Furthermore, compared to controls, the percentage of PCs that survived in P5-*PC-DTR* animals (∼66% of No DT controls) did not match the percentage of granule cells that were produced (∼81% of No DT controls). Thus, the ratio of PCs to granule cells that is important for proper CB circuitry formation is disrupted in P5-*PC-DTR* animals because granule cells are over-produced. These results reveal that independent of producing new PCs following their ablation, there are mechanisms of cell and organ size regulation that ensure recovery of CB size.

Finally, we tested whether P30 P1-*PC-DTR* or P5-*PC-DTR* mice recover motor function following recovery of PC numbers and CB size or only size, respectively. Remarkably P1-*PC-DTR* animals had no significant changes in their motor function compared to controls (Figure 4F-J). In contrast, compared to both No DT and P1-*PC-DTR* mice, P5-*PC-DTR* mice showed a significant reduction in their latency to fall from the rotarod and had a shorter stride (Figure 4F-G and I-J) with no change in grip strength (Figure 4H), demonstrating a motor behavior deficit. Thus, rapid production of new PCs by FEPs enables functional recovery following depletion of PCs at P1. Furthermore, achieving correct cell numbers and/or proportions are more important than maintaining CB size for functional recovery after injury in P5-*PC-DTR* mice.

In summary, we discovered a new regenerative process in the developing CB involving a previously unidentified and normally dormant progenitor population (FEPs) that acts as a developmental back-up population to buffer against early postnatal loss of postmitotic neurons. Proliferation of FEPs is stimulated by ablation of PCs at P1 and importantly the response is rapid (∼24 h), ensuring other components of the developing CB that are dependent on PCs for their expansion or differentiation are not compromised. However, FEPs decrease in number and capacity to regenerate during the first postnatal week, and consequently PCs are poorly replenished when ablated at P5. The cerebella of P5-*PC-DTR* mice nevertheless try to adapt by attaining near normal dimensions through a mechanism that includes increasing cell size (Figure4_supplement 5). The CB therefore has multiple mechanisms for regulating organ size following perinatal injury that depend on the precise stage of development. Furthermore, the motor deficits seen in P5-*PC-DTR* mice highlight the importance of maintaining the correct number of PCs during development, not just organ size.

The regenerative processes previously described in neuronal tissues involve adaptive reprograming of cells that are either actively proliferating or retain proliferative capacity and also have cell fate plasticity (Benner et al., 2013; Buffo et al., 2008; Jinnou et al., 2018; Lin et al., 2017; Llorens-Bobadilla et al., 2015; Lopez-Juarez et al., 2013; Marti-Fabregas et al., 2010; Robel et al., 2011; Samanta et al., 2015; Sirko et al., 2013; Wojcinski et al., 2017). Here we identified a distinct regenerative process that involves a local, unipotent and dormant progenitor. Unlike Nestin-expressing progenitors of the CB or astrocytes and neural stem cells in the forebrain that produce neurons upon injury, FEPs do not require reprograming and cell fate plasticity as they instead maintain their lineage and proliferate and mature upon injury. An important question raised by our study is whether regeneration of postmitotic neurons by age-dependent unipotent progenitors is unique to the CB, where protracted development might provide a conducive milieu, or whether all brain regions retain similar progenitors after each neuron subtype is generated. Furthermore, understanding the mechanisms of PC regeneration in newborn mice should provide insights that could enable regeneration in the adult brain.

## Acknowledgements

We thank past and present members of the Joyner laboratory for discussions and technical help. We thank T. Jessell and Jay Bikoff for providing the *FoxP2^Flpo^* line and P. Faust for sending us the *Pcp2^Cre^* line. We are grateful to M. E. Hatten, S. Shi, R. Sillitoe, A. Rosello-Diez and D. G. Placantonakis for comments on the manuscript. This work was supported by grants from the NIH to ALJ (R01NS092096 and R37MH085726) and a National Cancer Institute Cancer Center Support Grant [P30 CA008748-48].

## Materials and Methods

### Animals

All the experiments were performed according to protocols approved by the Memorial Sloan Kettering Cancer Center’s Institutional Animal Care and Use Committee (IACUC). Animals were given access to food and water *ad libitum*and were housed on a 12-hour light/dark cycle.

The following mouse lines were used for these experiments: *Pcp2^Cre^* (Zhang et al., 2004), *Nestin-CFP(Mignone, Kukekov, Chiang, Steindler, & Enikolopov, 2004; Wojcinski et al., 2017)*, *FoxP2^Flpo^*(Bikoff et al., 2016), *Rosa26^LSL-DTR^* (Stock no: 007900, The Jackson Laboratories)(Buch et al., 2005), *Rosa26^LSL-TdT^* (*ai14*, Stock no: 007909, The Jackson Laboratories)(Madisen et al., 2010), *Rosa26^FRT-STOP-FRT-TdT^* derived from *ai65* (Stock no: 021875, The Jackson Laboratories)(Madisen et al., 2015). Both sexes were used for analyses and no randomization was used. Exclusion criteria for experimental data points were sickness or death of animals during the testing period. No randomization was used and masking was used only for the behavior studies where the experimenter was blind to the genotypes.

Diphtheria toxin (30 µg/g of mouse; List Biological Laboratories, INC) was injected subcutaneously either at postnatal day (P) 1 or P5 and the brains were collected at various ages (Fig. 1a and Fig. 5a). Mice not given DT (No DT mice) were *Pcp2^Cre/+^; R26^DTR/LSL-TdT^* littermates and injected with the same volume of vehicle (PBS).

BrdU (50 µg/g of mouse; Sigma) was injected subcutaneously.

### Tissue Preparation and Histological Analysis

For P5 and younger animals, brains were dissected and fixed in 4% paraformaldehyde (PFA) for 24-48 hours (h) at 4°C. Animals older than P5 were anesthetized using intraperitoneal injection of a Ketamine (100 mg/kg) and Xylaxine (10 mg/kg) cocktail. Once full anesthesia was achieved, animals were systemically perfused with ice-cold PBS, followed by 4% PFA. Brains were dissected and post-fixed in 4% PFA for 24-48 h. Fixed brains were allowed to sink in 30% Sucrose in PBS solution and then embedded in OCT (Tissue-Tek) for cryosectioning. 14µm-thick cryosections were obtained using a Leica cryostat (CM3050S) and mounted on glass slides. Frozen sections were stored at −20°C for future analysis.

For immunofluorescent (IF) analysis, slides were allowed to warm to room temperature (RT). After washing once with PBS, slides were blocked using 5% Bovine Serum Albumin (BSA, Sigma) in PBS-T (PBS with 0.1% Triton-X) for 1 h at RT. Slides were then incubated overnight at 4°C with primary antibodies diluted in blocking buffer. **Figure1_source data 1.** summarizes the primary antibodies used in this study. Upon primary antibody incubation, slides were washed with PBS-T (3 × 5 minutes), incubated with specific AlexaFluor-conjugated secondary antibodies (1:500 in blocking buffer, Invitrogen) for 1 h at RT and then washed again with PBS-T (3 × 5 minutes). Counterstaining was performed using Hoechst 33258 (Invitrogen) and the slides were mounted with Fluoro-Gel mounting media (Electron Microscopy Sciences).

Haematoxylene and Eosin (H&E) staining was performed to assess cerebellar cytoarchitecture and measure area (size).

### Image Acquisition and Analysis

Images were collected either with a DM6000 Leica microscope or Zeiss LSM 880 confocal microscope and processed using ImageJ Software (NIH).

For each quantification, 3 midline parasagittal sections/brain were analyzed and data was averaged. Cells were counted using the Cell Counter plugin for ImageJ (NIH). Analyses of PC and FEPs numbers were performed by counting all of the PCs on a midline parasagittal section. CB area was calculated by defining a region of interest by outlining the perimeter of the outer edges of the CB, using ImageJ. EGL thickness was calculated by dividing the area of the EGL by the length of the EGL in midline sections. IGL density was calculated by counting the number of nuclei in three 40x fields from lobule 8 in three midline sections and by dividing the number by the area of the region counted. PC soma size and dendrite length were calculated using randomly distributed TdT+ PCs from 3 midline sections (>20 cells/section). Soma area is calculated by outlining the perimeter of the outer edges of each cell. Cells that show primary dendrites were used for this analysis to ensure that the region where the maximum soma area is observed is used for the analyses. For dendrite length quantifications, primary and secondary dendrite length was measured and summed and PCs around the base of fissures were omitted.

### Behavioral Testing

5-week old animals (No DT: n = 17, DT@P1: n = 9 and DT@P5: n = 11) were used to assess differences in motor behavior. The same sets of mice were used for all three tests described below.

#### Rotarod

An accelerating rotarod (47650; Ugo Basile) was used for these experiments. Animals were put on the rod, and allowed to run till the speed reached to 5 rpm. Then the rod was accelerated from 5 to 40 rpm over the course of 300 seconds. Recording was stopped at 305 seconds. Time of fall was recorded for each animal. Analysis was performed 3 times a day on 3 consecutive days. Between each trial, animals were allowed to rest for 10 minutes in their home cage.

#### Grip Strength

To test whether any effects observed in the rotarod test were due to muscle weakness, grip strength measurements were performed using a force gauge (1027SM Grip Strength Meter with Single Sensor, Columbus Instruments). Animals were allowed to hold a horizontal grip while being gently pulled away by the base of their tail. Measurements were performed 5 times with 5 minute resting periods in between. Force amount was recorded. Data was normalized to mouse’s weight and represented in (Force/gram).

#### Footprinting Analysis

Forefeet and hindfeet were painted with red and blue nontoxic acrylic paint (Crayola), respectively. Animals were allowed to walk on a strip of paper laid along the floor of a 50 cm long, 10 cm wide custom-made Plexiglas tunnel with a dark box at the far end. 3 runs/mouse were performed and the distances between the markings were measured.

### Statistical Analysis

Prism (GraphPad) was used for all statistical analysis. Statistical comparisons used in this study were Student’s two-tailed t-test; One-way and Two-way analysis of variance (ANOVA), followed by post hoc analysis with Tukey’s test for multiple comparisons. Relevant F-statistics and p-values are stated in the figure legends and the p-values of the relevant post hoc multiple comparisons are shown in the figures. Summary of all the statistical analysis performed can be found in **Figure 1_source data 2.** The statistical significance cutoff was set at P < 0.05. Population statistics were represented as mean ± standard deviation (SD) of the mean. No statistical methods were used to predetermine the sample size, but our sample sizes are similar to those generally employed in the field. n ≥ 3 mice were used for each experiment and the numbers for each experiment are stated in the figure legends.

## Supplementary Figures and Tables

**Figure 1_Supplement 1.**
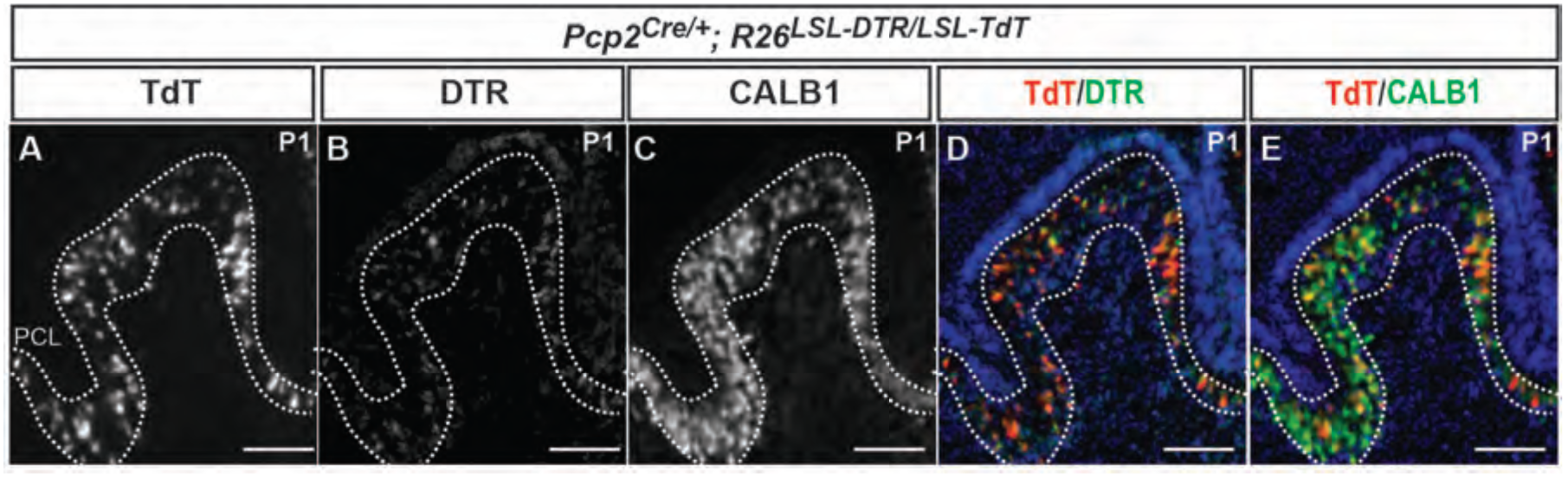
Characterization of DTR and TdT expression in PCs of *PC-DTR* mice at P1. **A-E**. IF analysis of **(A)** TdT, **(B)** DTR, **(C)** CALB1 and combination shows that all the TdT+ cells express DTR **(D)** and CALB1 **(E)**. DTR: Diphtheria toxin receptor, PCL: Purkinje cell layer. Scale bar: 100 µm

**Figure 1_Supplement 2.**
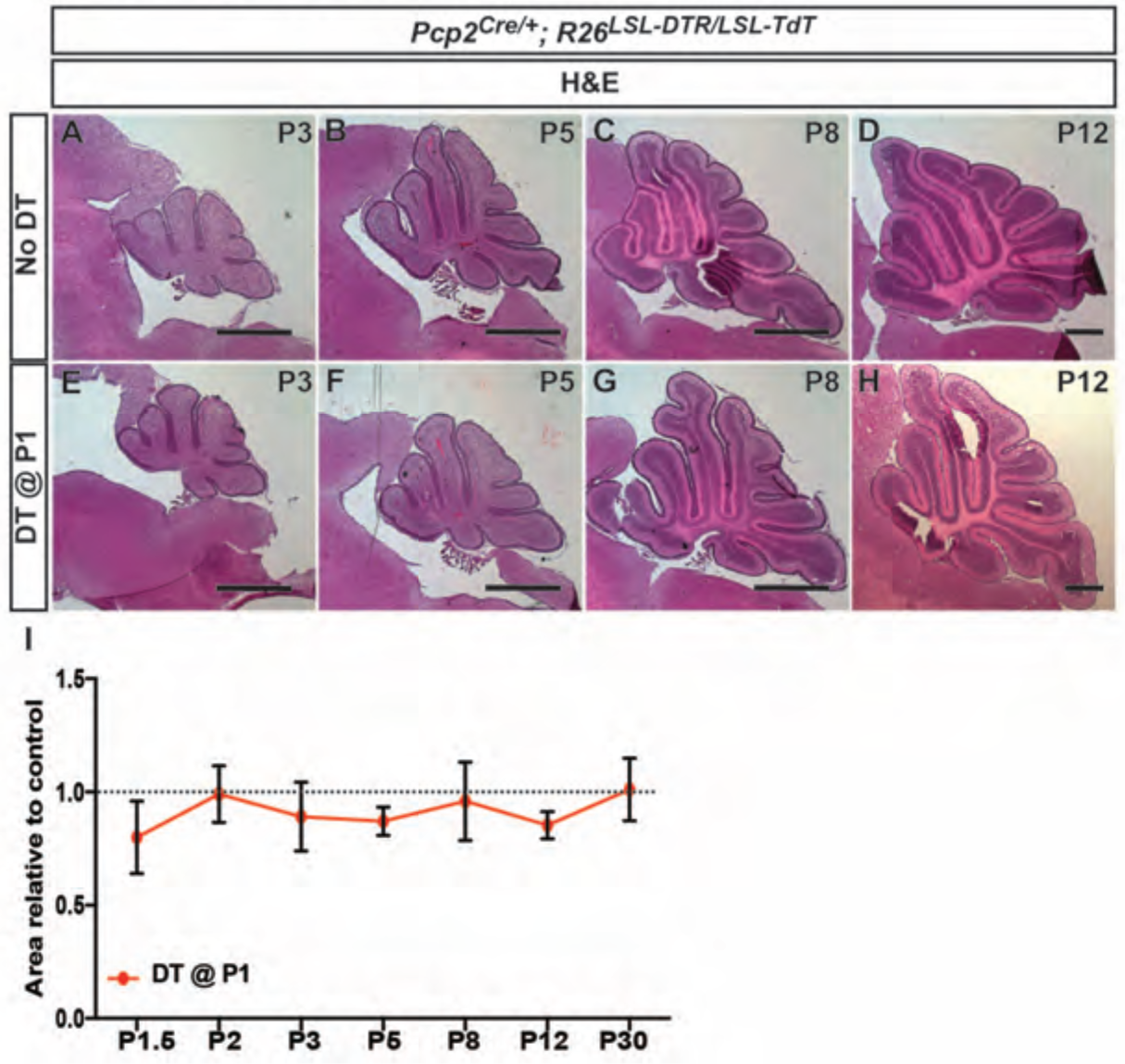
CB size and morphology appears normal following DT-mediated ablation of PCs at P1. **A-H**. H&E stained midline sagittal sections of cerebella at the ages indicated for No DT (**A-D**) and P1-*P**C-D**TR* (**E-H**) mice. **I**. Quantification of midline sagittal areas of cerebella shows no differences upon DT injection (n ≥ 3 for each age). Scale bars: 500 µm

**Figure 1_Supplement 3.**
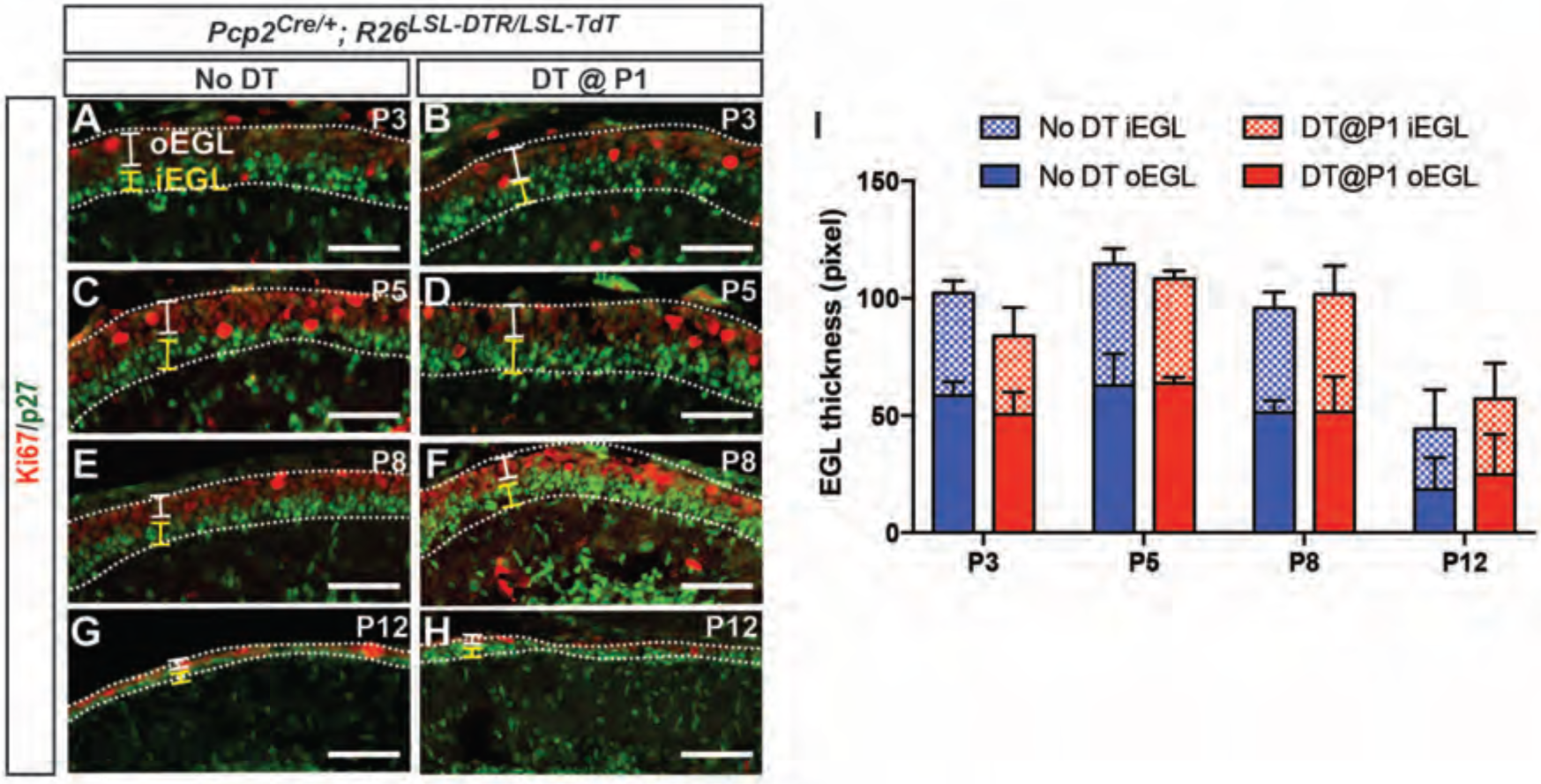
External granule cell layer thickness is not changed after DT-mediated killing of PCs at P1. **A-H**. IF analysis of Ki67 (outer EGL, oEGL) and p27 (inner EGL, iEGL) in No DT (A, C, E, G) and P1-*PC-DTR* (B, D, F, H) animals at the indicated ages. **I.** Quantification of the thickness (area/length) of the oEGL, which contains proliferating granule cell progenitors, and the iEGL, which contains the differentiating granule cells, reveals no significant differences in total EGL area and the ratio of inner and outer EGL areas between No DT and P1-*P**C-D**TR* animals (n = 3/condition) (P = 0.85). EGL: external granule layer. Scale bars: 100 µm

**Figure 2_Supplement 1.**
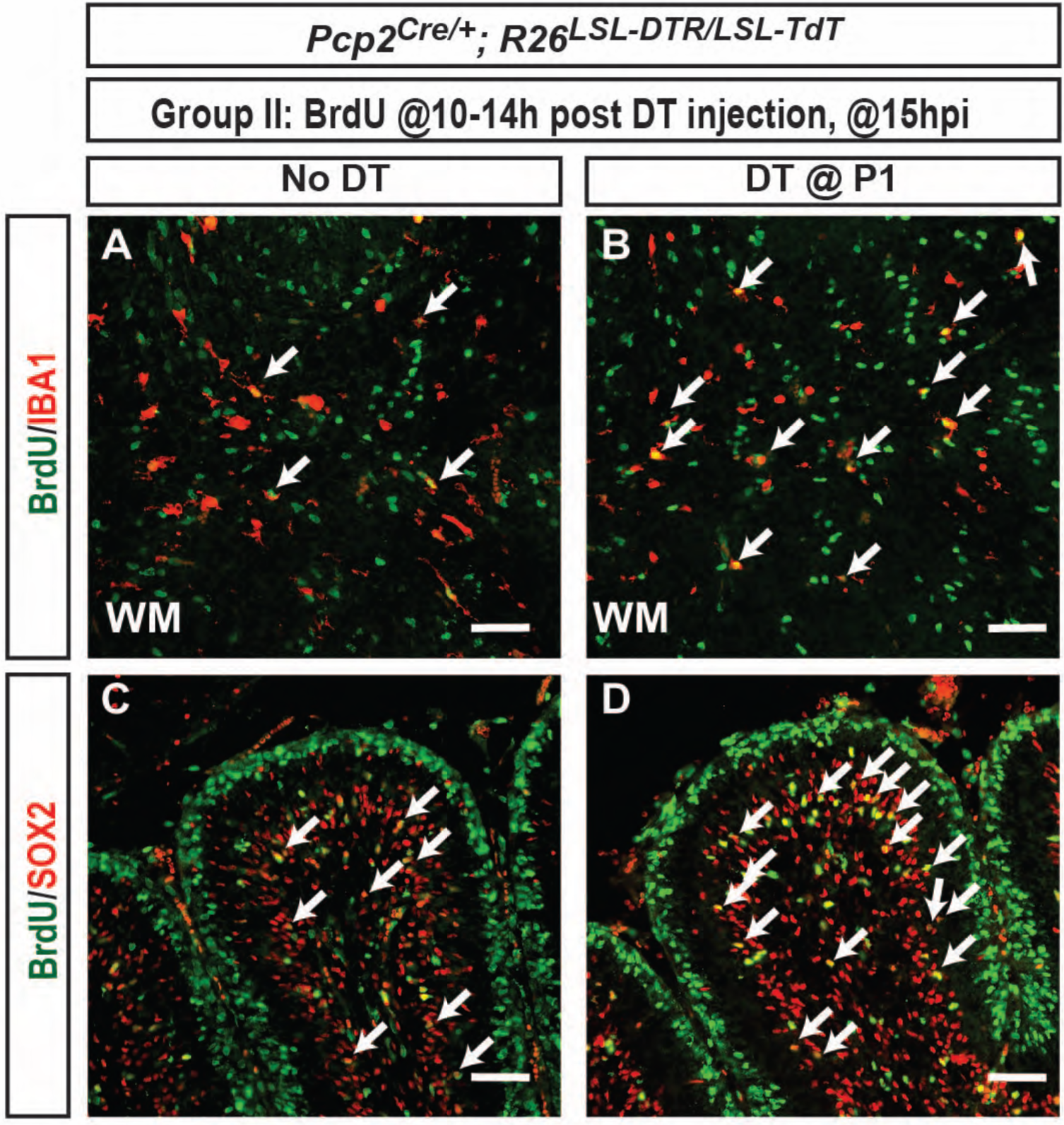
Characterization of proliferating cell types that respond to DT-mediated ablation of PCs. **A-D**. IF analysis of BrdU+ cells indicates that number of proliferating (**A-B**) IBA1+ microglia and (**C-D**) SOX2+ cells (astrocytes and NEPs) increased upon ablation of PCs in P1-*P**C-D**TR* mice. Arrows show BrdU+ IBA1+ and Sox2+ cells. Scale bars: 100 µm

**Figure 2_Supplement 2.**
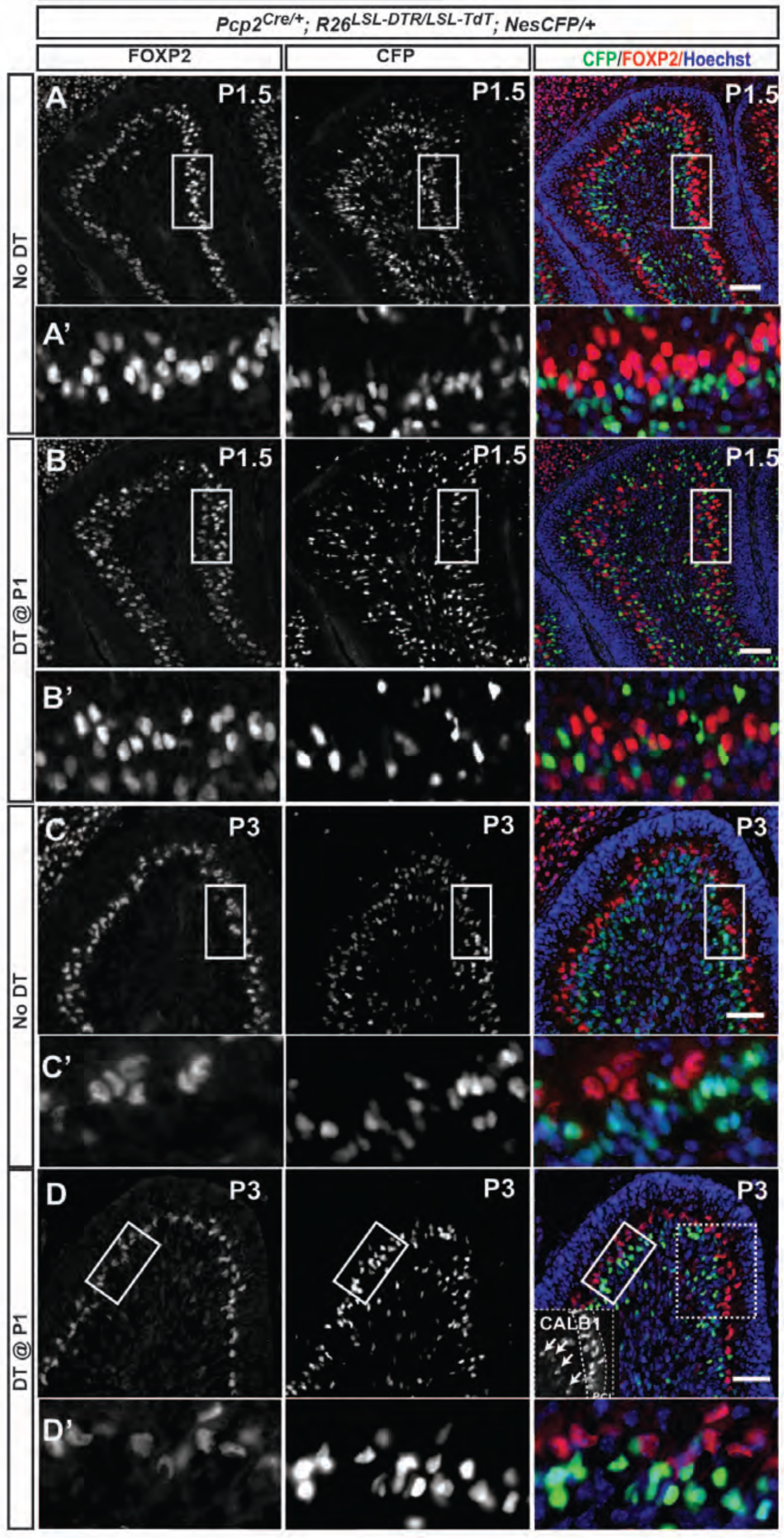
NEPs are not responsible for the recovery of PCs following DT-mediated ablation at P1. **A-D**. A *Nestin-CFP* reporter was used to transiently track the fate of NEPs and revealed no overlap between FOXP2+ cells and CFP staining 12 h (P1.5) (A, A’ and B, B’) and 2 days (P3) (C, C’ and D, D’) after PC depletion in P1-*P**C-D**TR* mice. Note that in B’ FOXP2 staining is weaker in the ectopic layer of dying PCs that is forming on the inside of the PCL. Inset in d shows the ectopic CALB1+ cells. NEP: Nestin-expressing progenitors. Scale bars: 100 µm

**Figure 3_Supplement 1.**
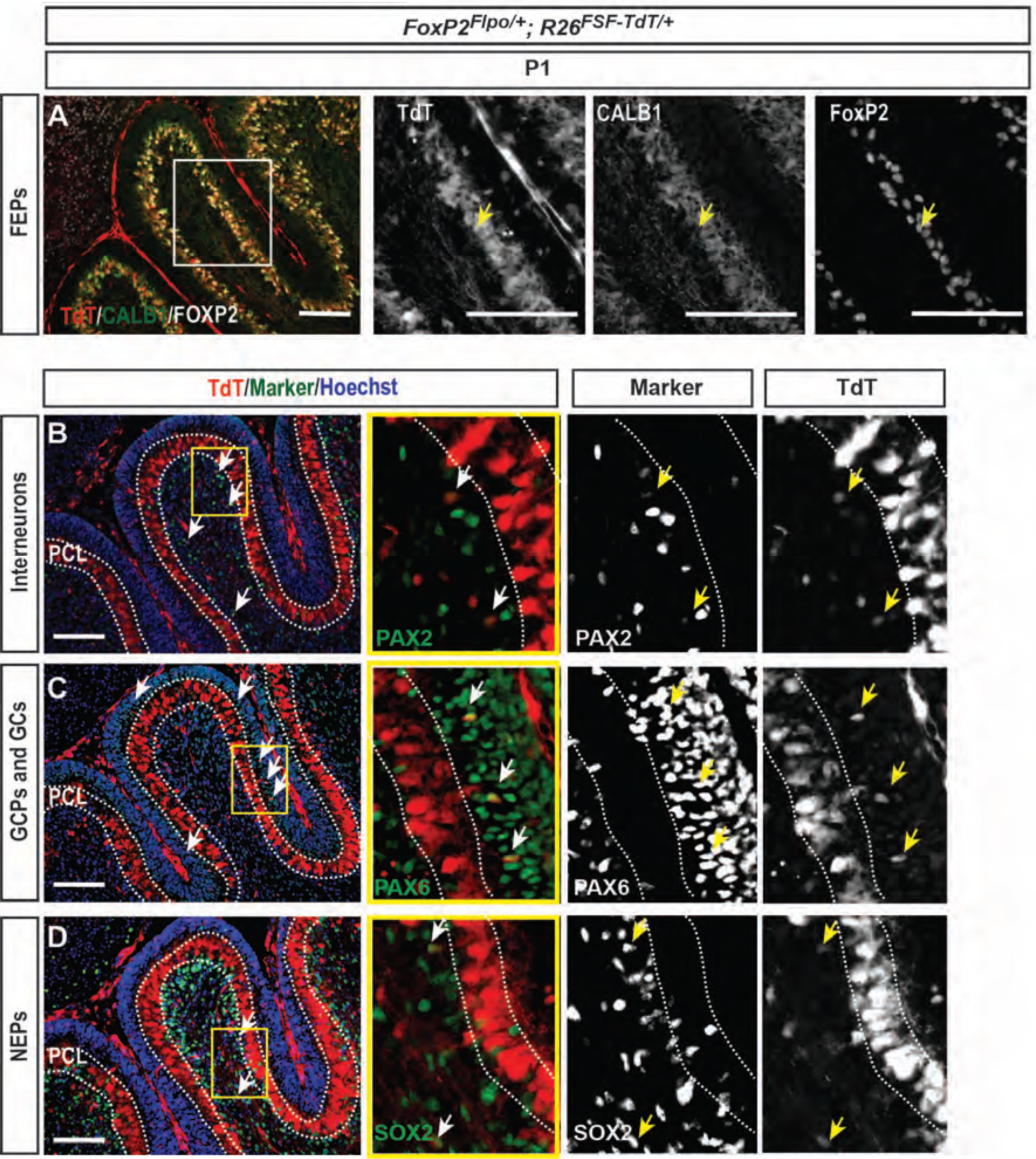
*FoxP2-TdT* fate mapping at P1 marks FEPs in the PCL as well as other rare cells outside of PCL. *FoxP2^Flpo/+^; R26^FSF-TdT/+^* (*FoxP2-TdT*; FSF = frt-stop-frt) animals were analyzed at P1. **A**. As predicted, all of the FOXP2+ cells in the PCL were labeled with TdT+ and some were CALB1-/low. Arrow shows a TdT+, FOXP2+ CALB1-/low cell (FEP) in the PCL. **B-D**. *FoxP2-TdT* also marks rare Pax2+ interneurons **(B)**, Pax6+ granule cells **(C)** and Sox2+ glial cells/progenitors **(D)**, none of which reside in the PCL. These results suggest that *Flpo* allele is unexpectedly expressed transiently in rare embryonic progenitors of other lineages than PCs. Scale bars: 200 µm

**Figure 3_Supplement 2.**
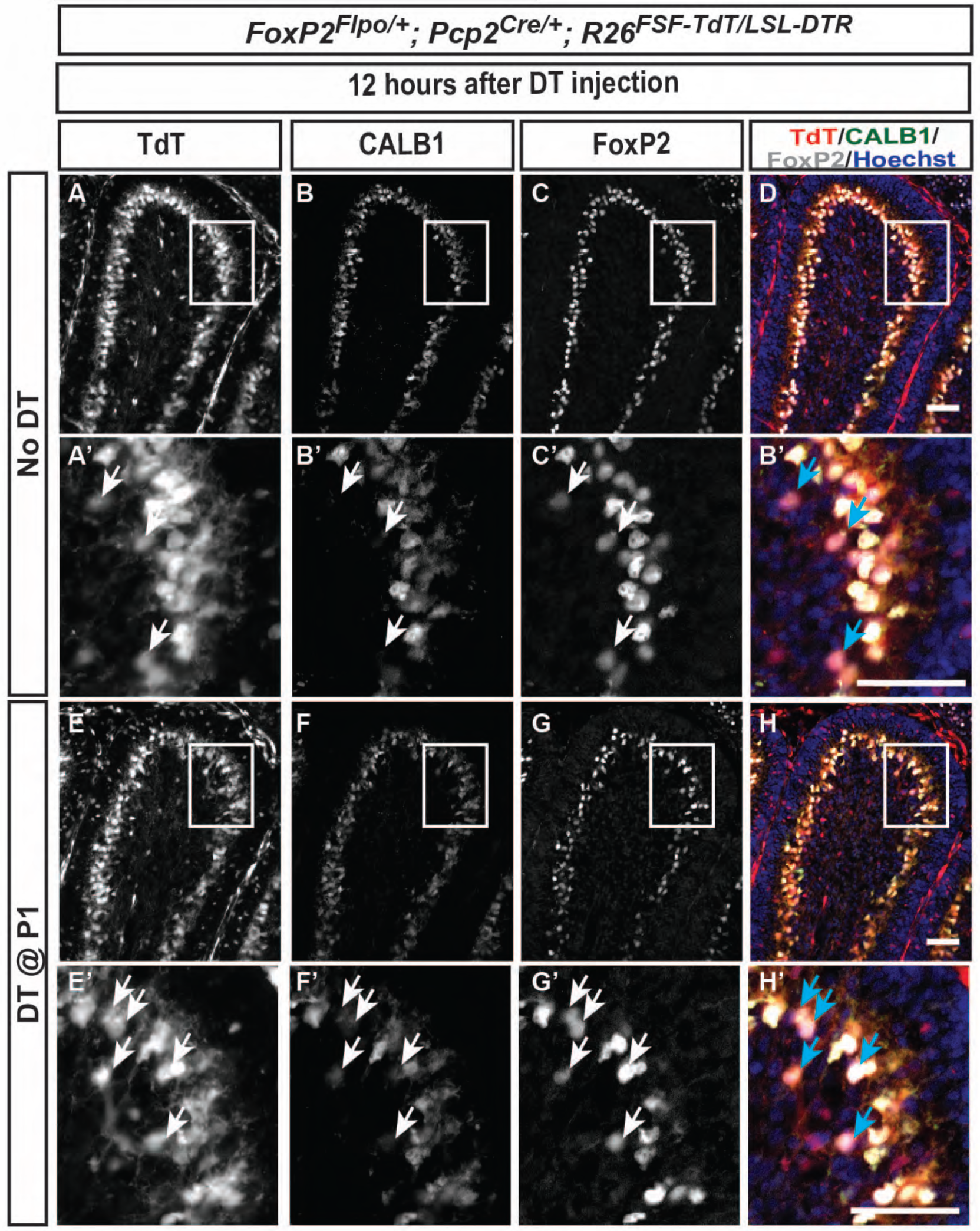
The number of *FoxP2-TdT*-marked FEPs increases 12 hours after DT injection at P1. **A-H**. FEPs (TdT+, FOXP2+, CALB1-/low, arrows in the higher magnified images) are sparsely located in No DT *FoxP2-TdT* pups (**A-F**) and the number increases 12 hours after DT injection at P1 (**E-H**). Scale bars: 50 µm

**Figure 3_Supplement 3.**
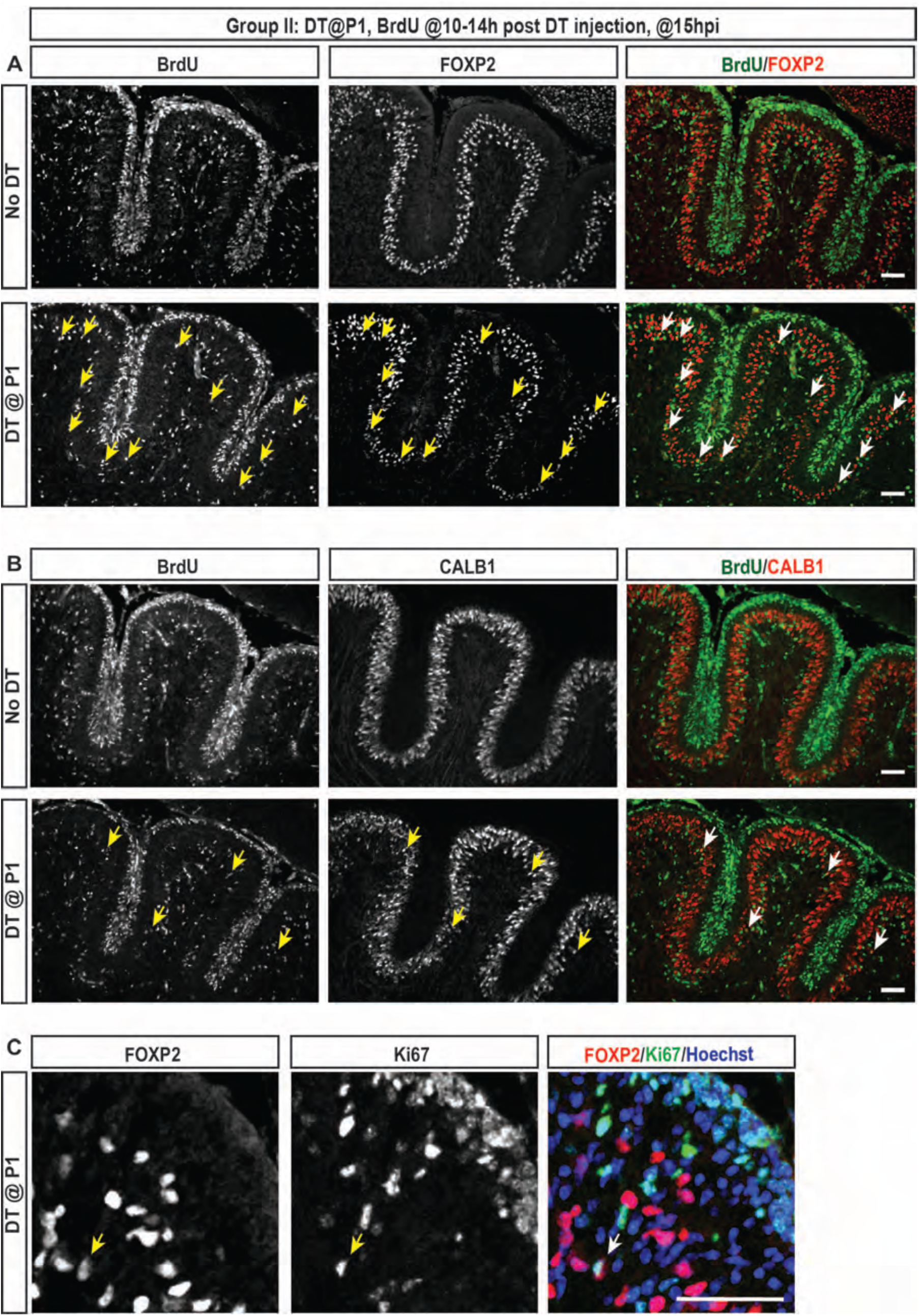
IF analysis of PCs at P1.5 (15 h post DT injection at P1) shows that FoxP2+ cells proliferate and there are more FoxP2+ cells that incorporate BrdU, compared to CALB1+ cells that are BrdU+. **A-B**. Analysis of co-labeling of FoxP2 **(A)** or CALB1 **(B)** with BrdU (injected 10-14 h post DT) at 15 hpi of DT shows that more FoxP2+ cells incorporate BrdU upon DT injection (lower panels) of P1-*P**C-D**TR* mice, compared to FoxP2+ CALB1+ cells. Brains of No DT mice did not show any PCs that incorporated BrdU (top panels). **C.** Confocal microscopy reveals FoxP2+ Ki67+ cells 15 hours post DT injection, further confirming proliferation of FOXP2+ cells in P1*-P**C-D**TR*. Arrows indicate BrdU or Ki67+ PCs that are either FoxP2+ or CALB1+. Scale bars: 100µm

**Figure 4_Supplement 1.**
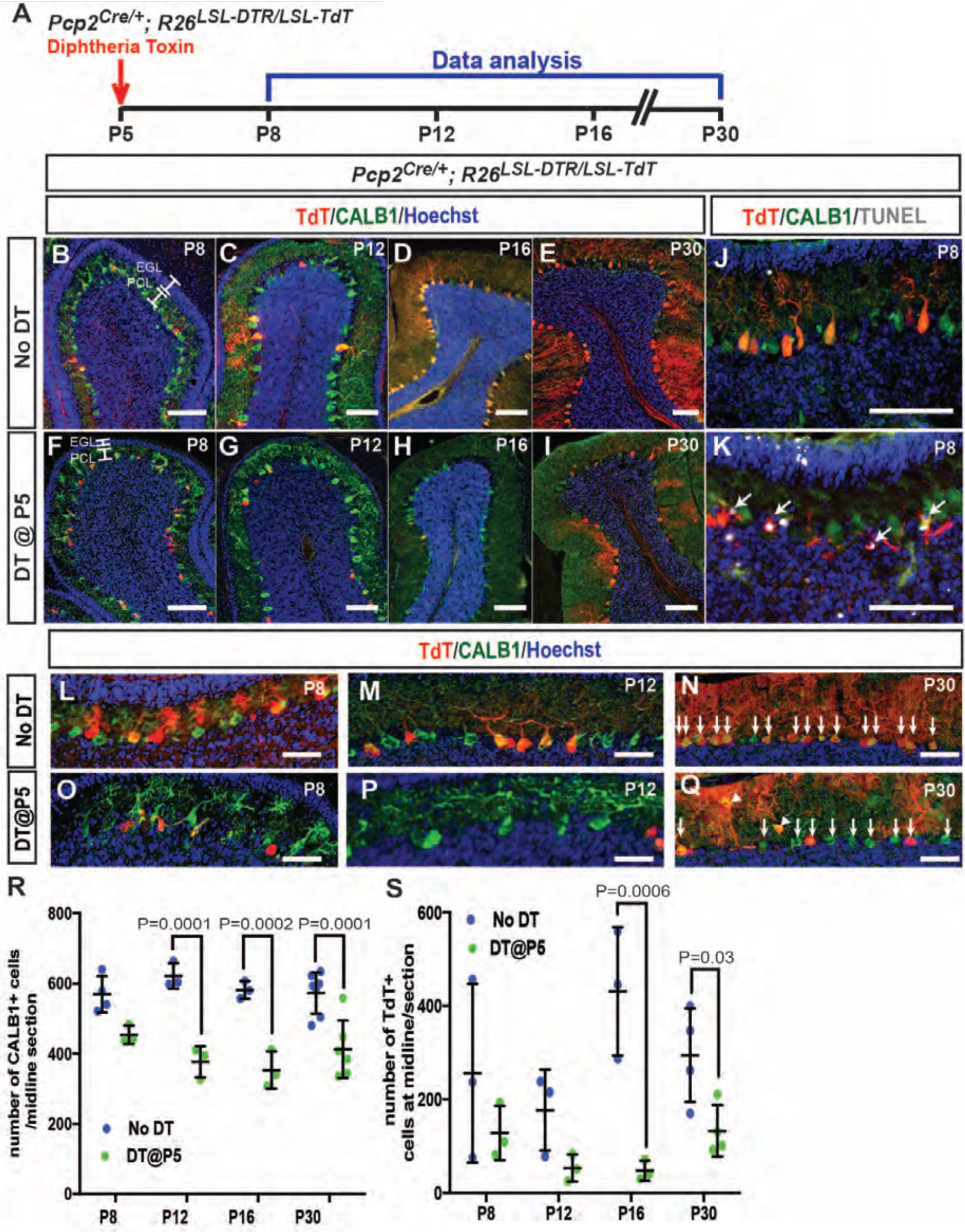
PC numbers are reduced upon PC ablation at P5-*PC-DTR*. A. Schematic representation of the experimental plan. **B-I**. IF analysis of PCs upon ablation at P5 (F, G, H, I) reveals lack of recovery of PC numbers. **J-K**. Analysis of apoptosis by TUNEL reveals TUNEL+ TdT cells (arrows) in PCL of P5-*PC-DTR* mice **(K)** but not is No DT mice **(J)** at P8**. **L-Q**.** Higher magnification of PCs from P8, P12 and P30 P5-P**C-D**TR animals and No DT controls (L, M, N) reveal that P5-*PC-DTR* mice (O, P, Q) have disrupted PC morphology and reduction in their numbers. Arrows show PCs and reduction in their number at P5-*PC-DTR* mice. Arrowheads: Ectopic PCs. **R.** Quantification of CALB1+ cells shows that PC numbers do not recover from ablation of PCs at P5 (Two-way ANOVA, F_(1,24)_=77.85, P = 0.0001, n ≥ 3). **S.** Quantification of the number of TdT+ cells, shows a large variation in recombination efficiency in No DT brains, and an initial decrease in TdT+ cells after DT injection at P5 (Two-way ANOVA, F_(1,18)_=26.29, P = 0.0001, n ≥ 3). At P30, P5-*PC-DTR* brains show decrease in the number of TdT+ cells compared to No DT animals (t-test, P = 0.03, n ≥ 4), similar to P1-*PC-DTR* animals (Fig. 1q). Significant *post hoc* comparisons are shown in the figure. EGL: External granule layer, PCL: Purkinje cell layer. Scale bars: a-k: 100 µm, l-q: 50 µm

**Figure 4_Supplement 2.**
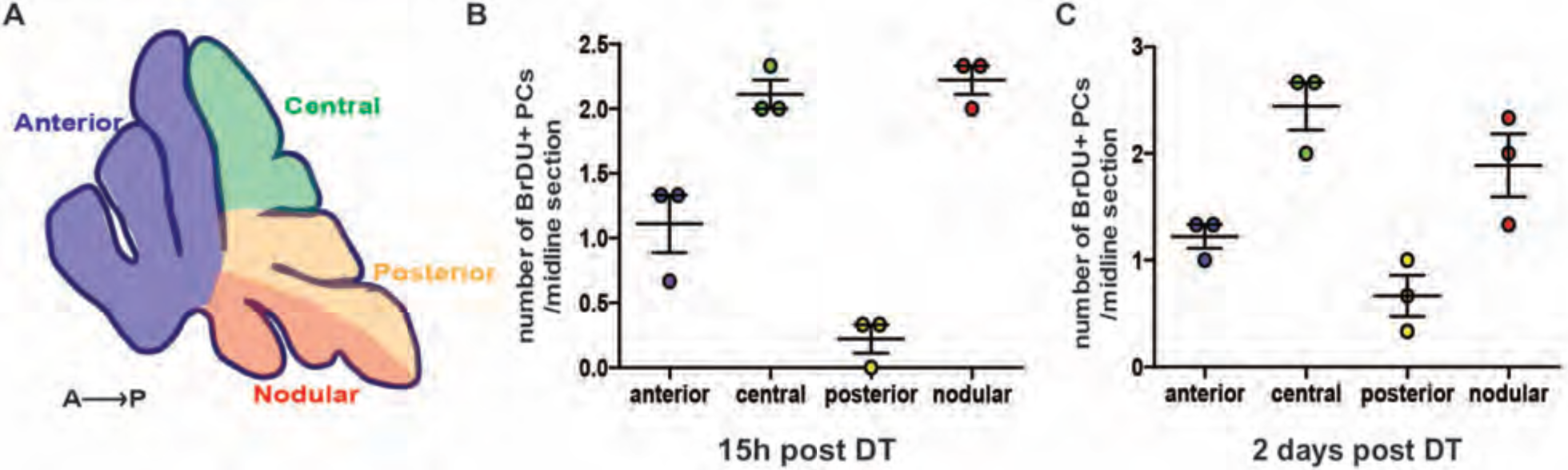
Distribution of BrDU+ PCs in P5-*P**C-D**TR* mice at 15 hours and 2 days post injection of DT. A. Schematic showing the different zones of the CB in a P5 sagittal midline section. **B-C**. Distribution of BrdU+ PCs across different zones analyzed 15 h **(B)** and 2 days **(C)** in P5-*P**C-D**TR* animals reveals that incorporation of BrdU is limited, and most of the cells reside in the central and the nodular zones, correlating with the localization of FEPs (n = 3 /condition). No BrdU incorporation was detected in No DT mice at the same ages.

**Figure 4_Supplement 3.**
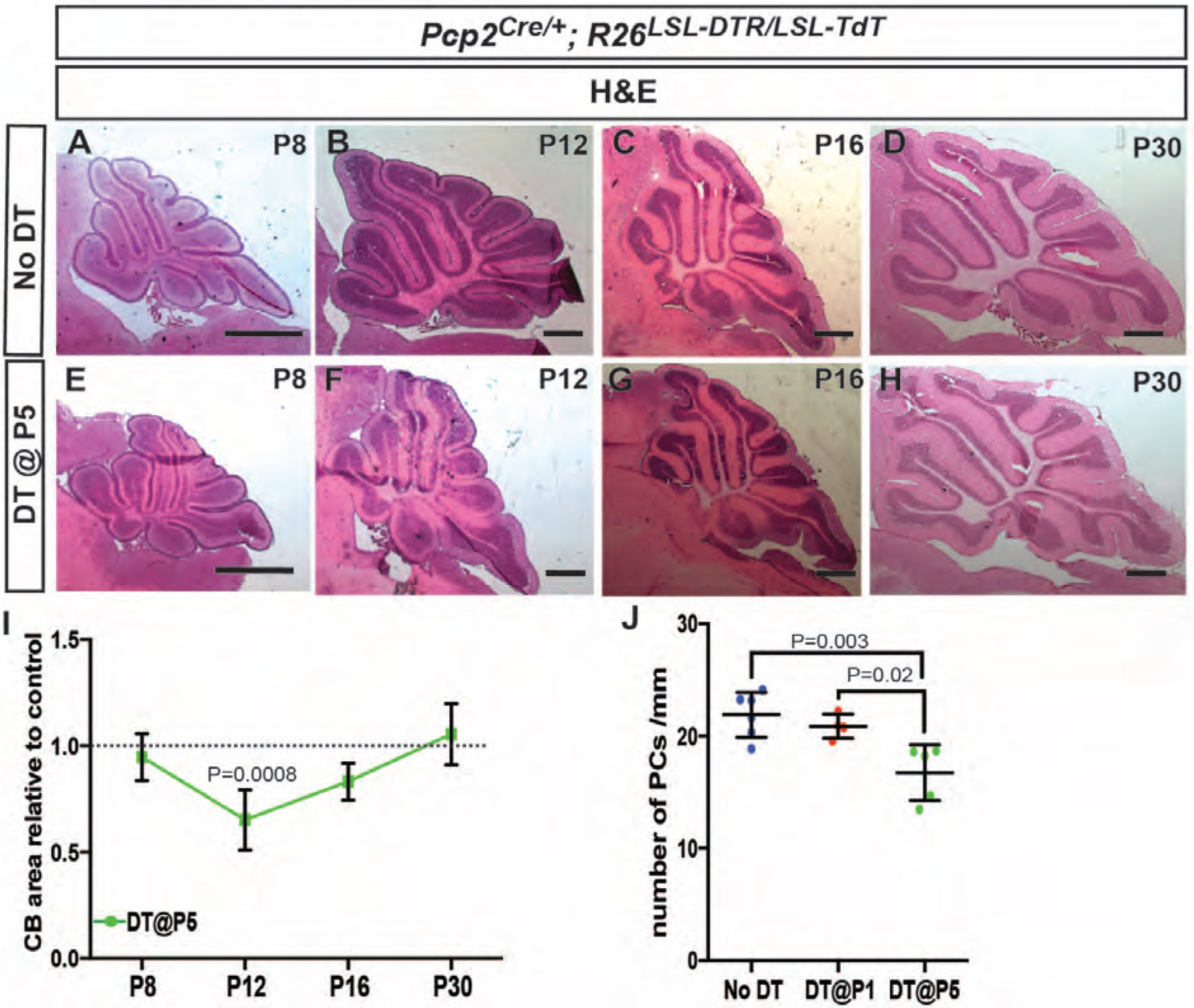
Transient decrease in CB size and altered PC characteristics after ablation of PCs at P5. **A-H**. H&E staining shows that the area of sagittal CB sections is reduced at P12 after DT injection in P5-*P**C-D**TR* mice No DT (F compared to B) and the significant difference is lost at P16 and P30. **I.** Quantification of CB area in midline sagittal sections demonstrates that CB size is smaller at P12. (Two-way ANOVA, F_(1,22)_=7.799, P = 0.01, n ≥ 3) **J.** The density of PCs was reduced at P30 in P5-*P**C-D**TR* but not in P1-*P**C-D**TR* animals, correlating with lack of recovery of PC numbers in parallel with recovery of CB area in the former mice (One-way ANOVA, F_(2,12)_=9.687, P = 0.003, n ≥ 4). Significant *post hoc* comparisons are shown in the figure. Scale bars: 500 µm

**Figure 4_Supplement 4.**
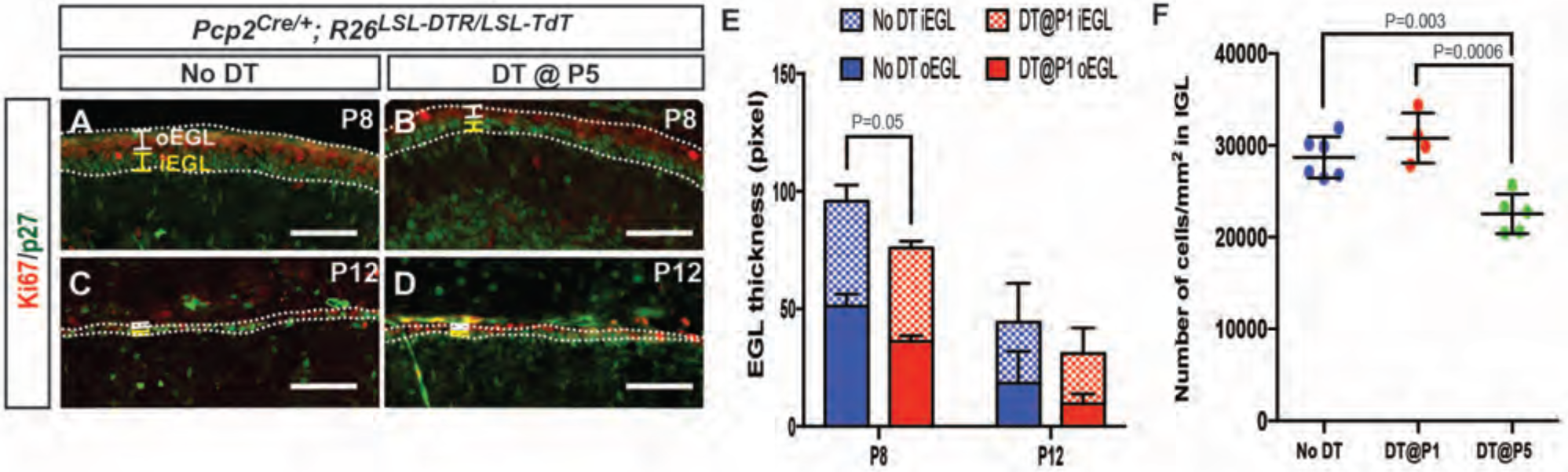
Analysis of external granule cell layer thickness after DT injection at P5. **A-D**. IF analysis of Ki67 (outer EGL, oEGL) and p27 (inner EGL, iEGL) in No DT (A, C) and P5-*P**C-D**TR* (B, D) mice. **E.** Quantification shows that both the oEGL and iEGL thicknesses (area/length) were significantly reduced at P8 (Two-tailed t-test, P = 0.05, n = 3), but not at P12**. F.** Likely as a consequence of a thinner EGL, granule cell density in the internal granule cell layer (IGL) is reduced only in P5-*P**C-D**TR* animals, but not in No DT and P1-*PC-DTR* animals (One-way ANOVA, F_(2,12)_=15.73, P = 0.0004, n ≥ 4). Significant *post hoc* comparisons are shown in the figure. Scale bars: 100 µm

**Figure 4_Supplement 5.**
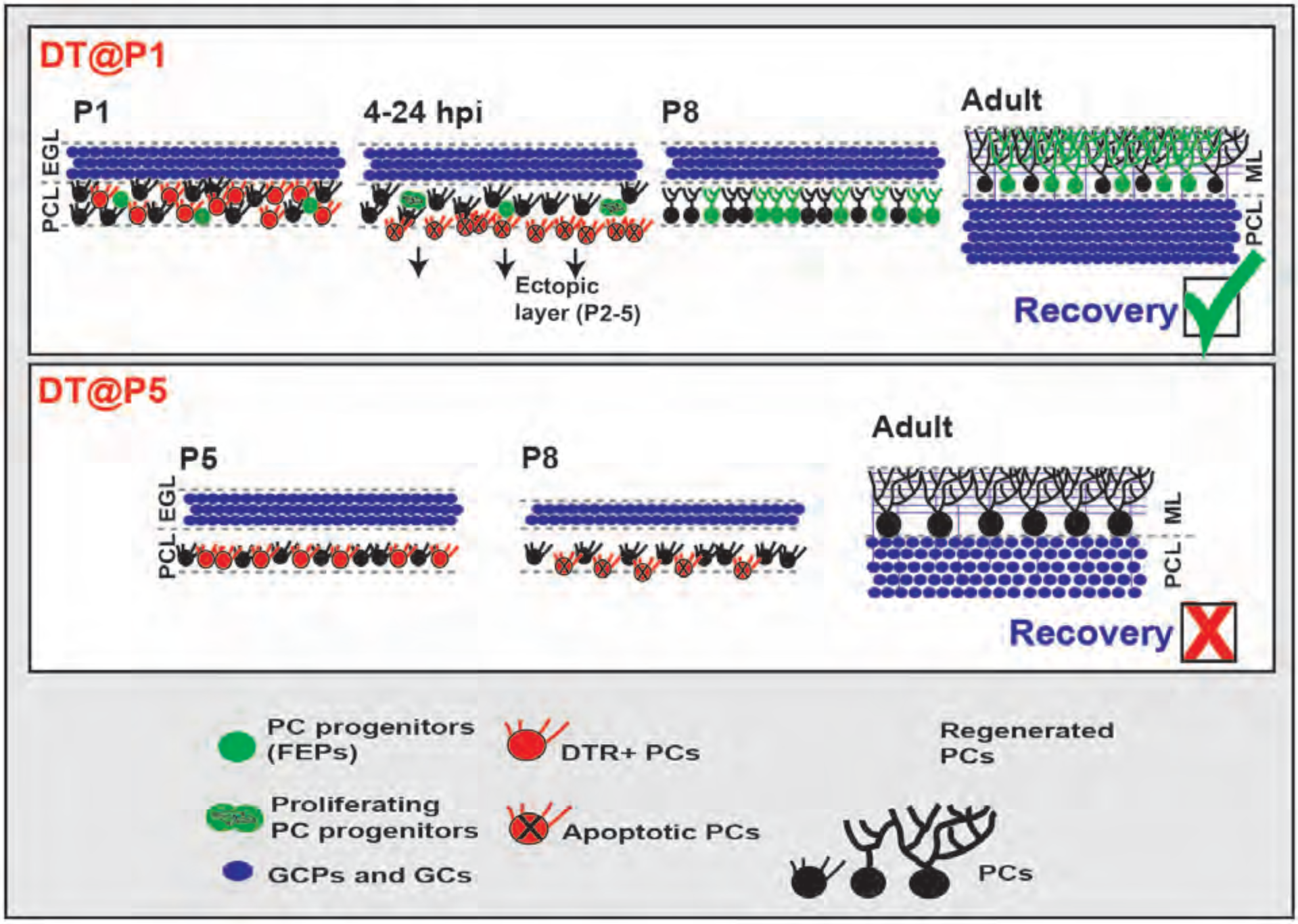
Graphical summary of the findings. FEPs: FoxP2-expressing progenitors, EGL: external granule cell layer, PCL: Purkinje cell layer, ML: Molecular Layer, GCP: granule cell progenitors, GC: granule cells.

**Figure 1_source data 1.**
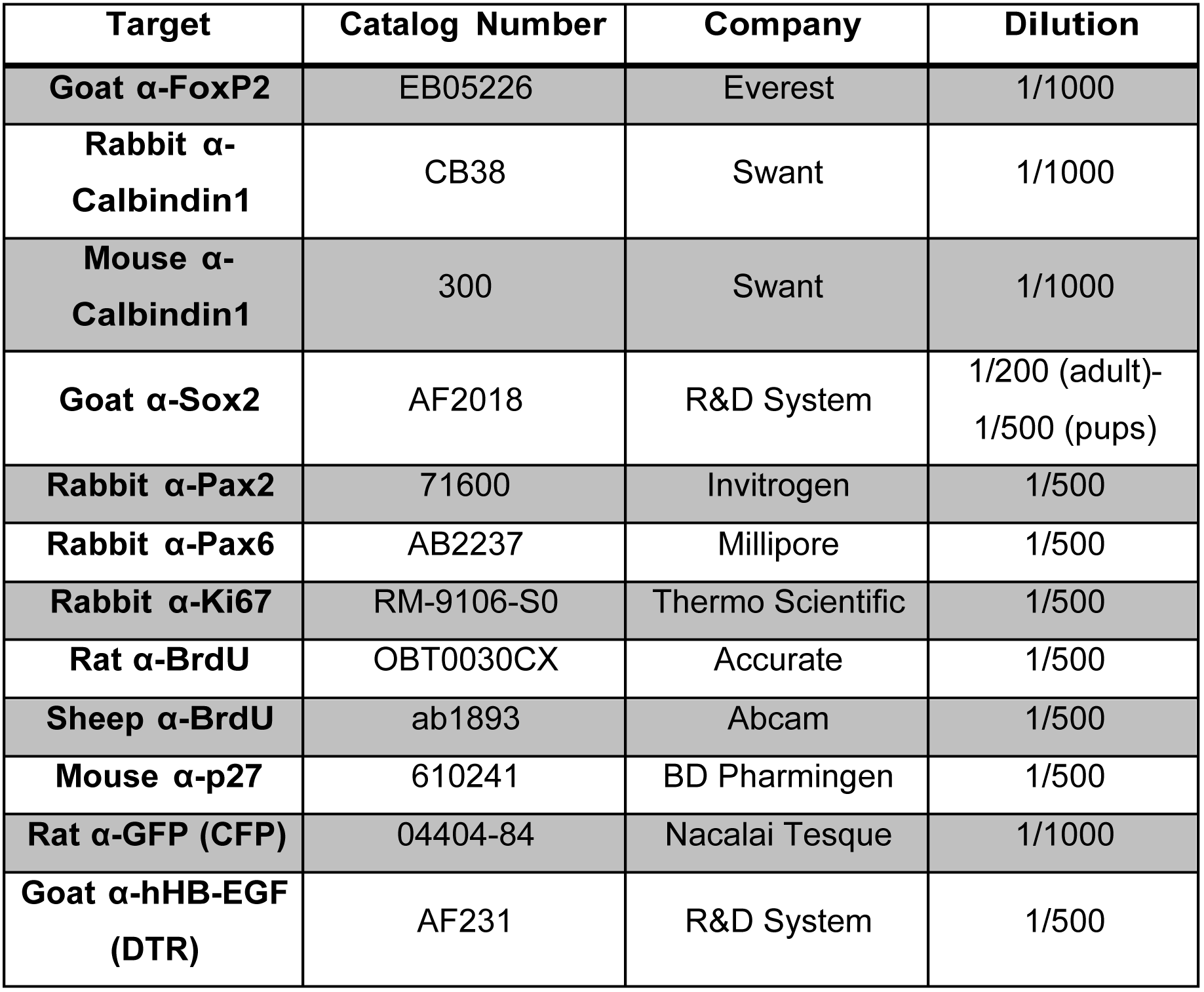
Summary of the antibodies used in the study.

**Figure 1_source data 2.**
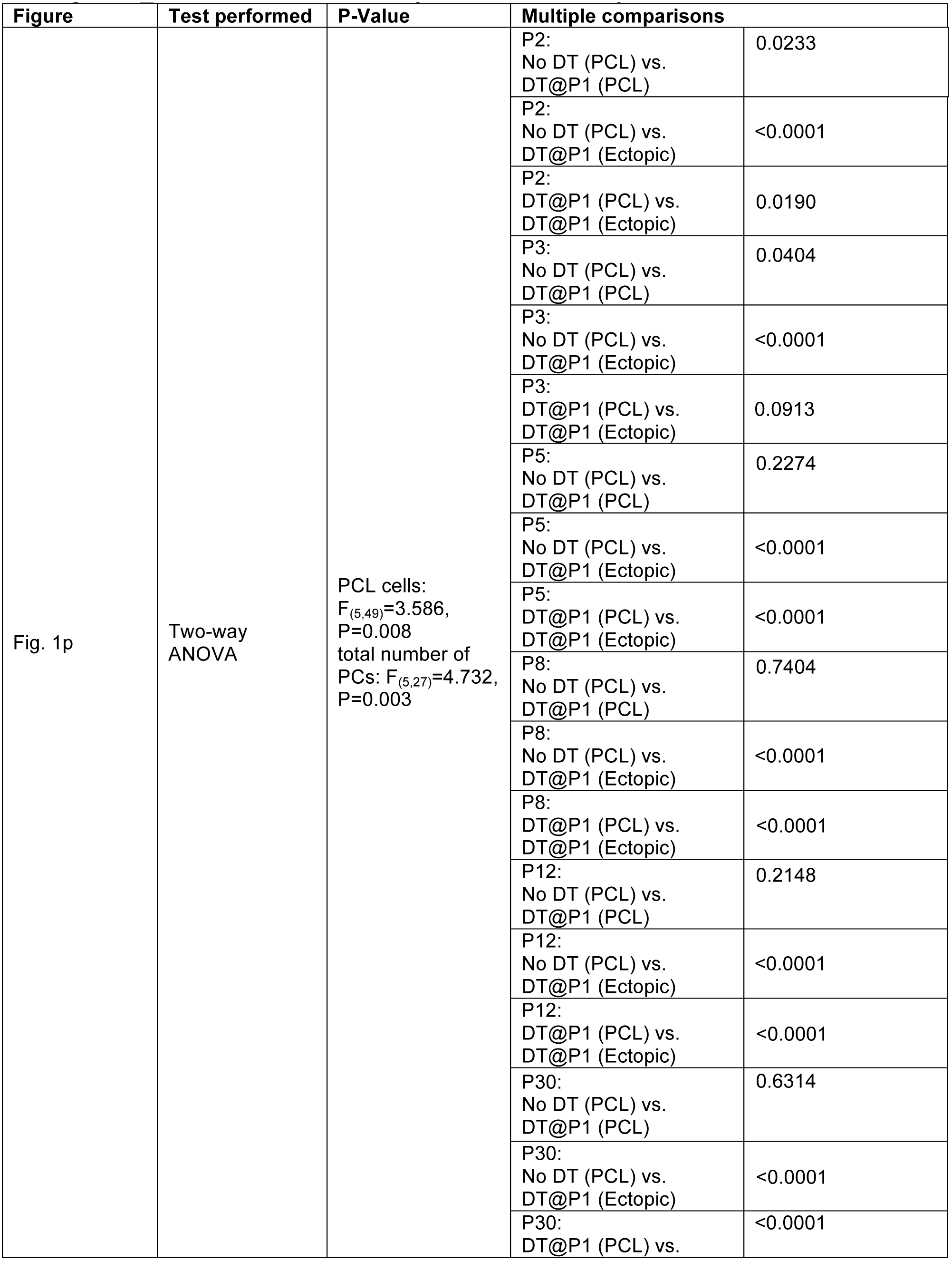
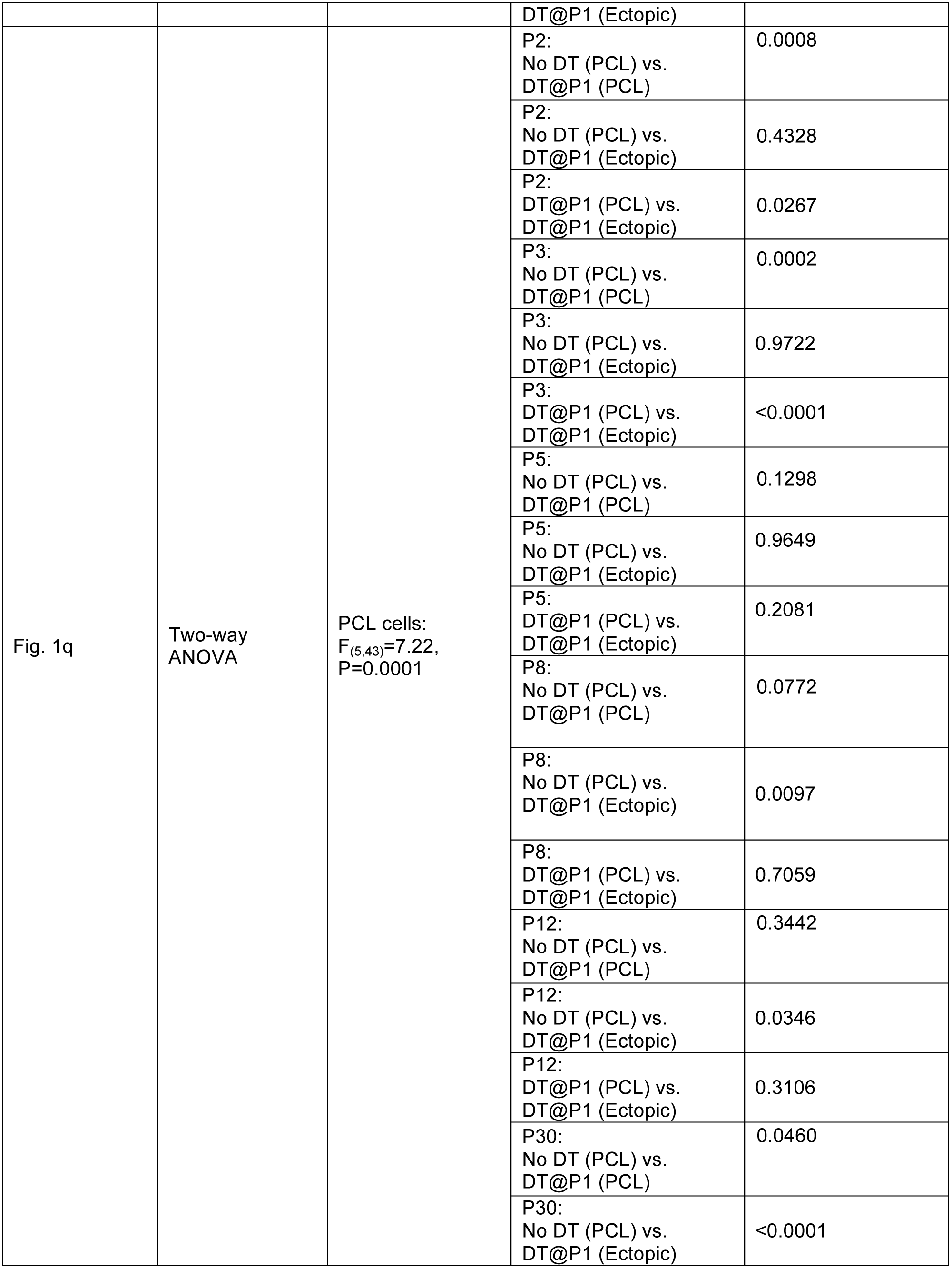
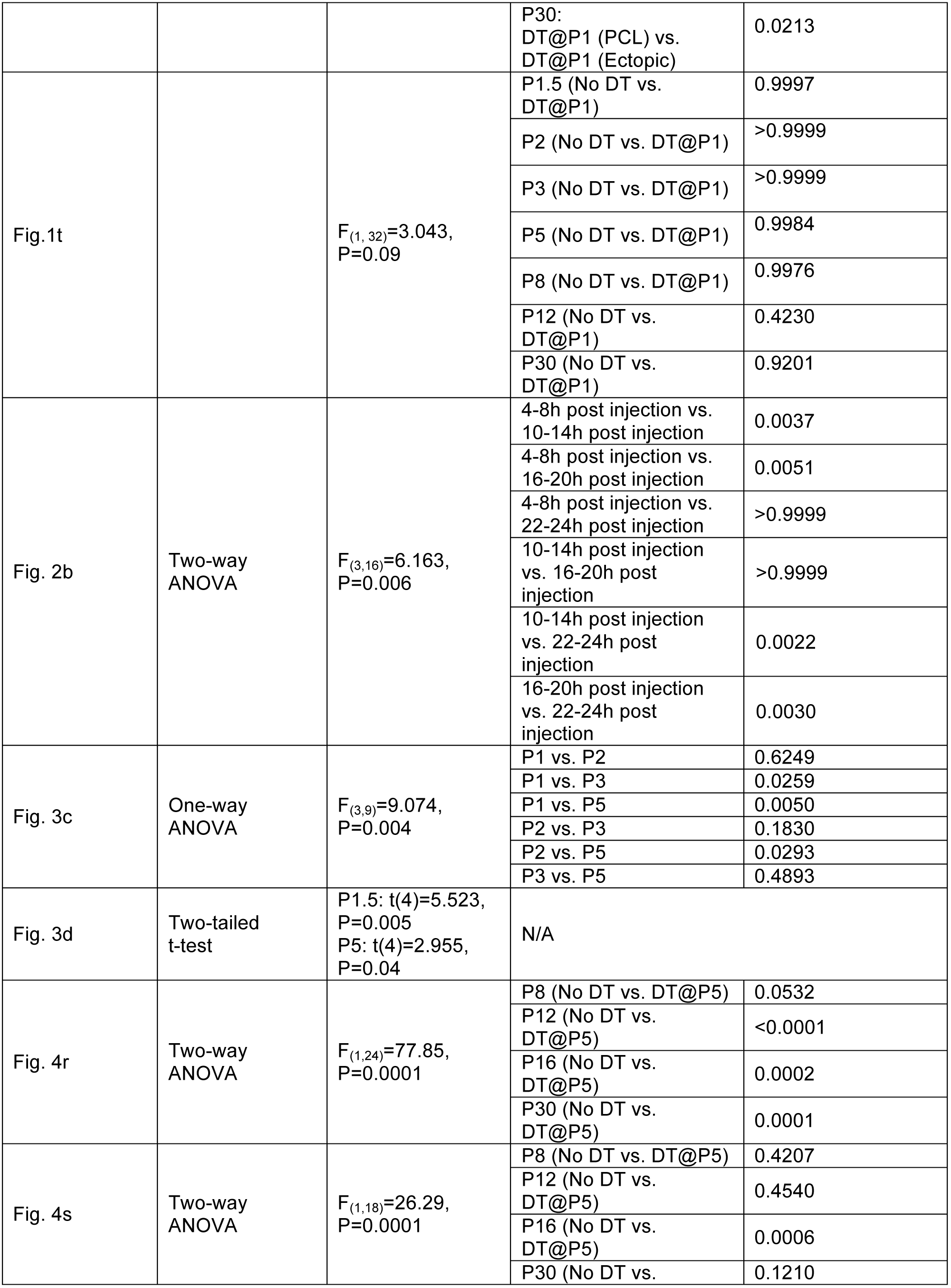
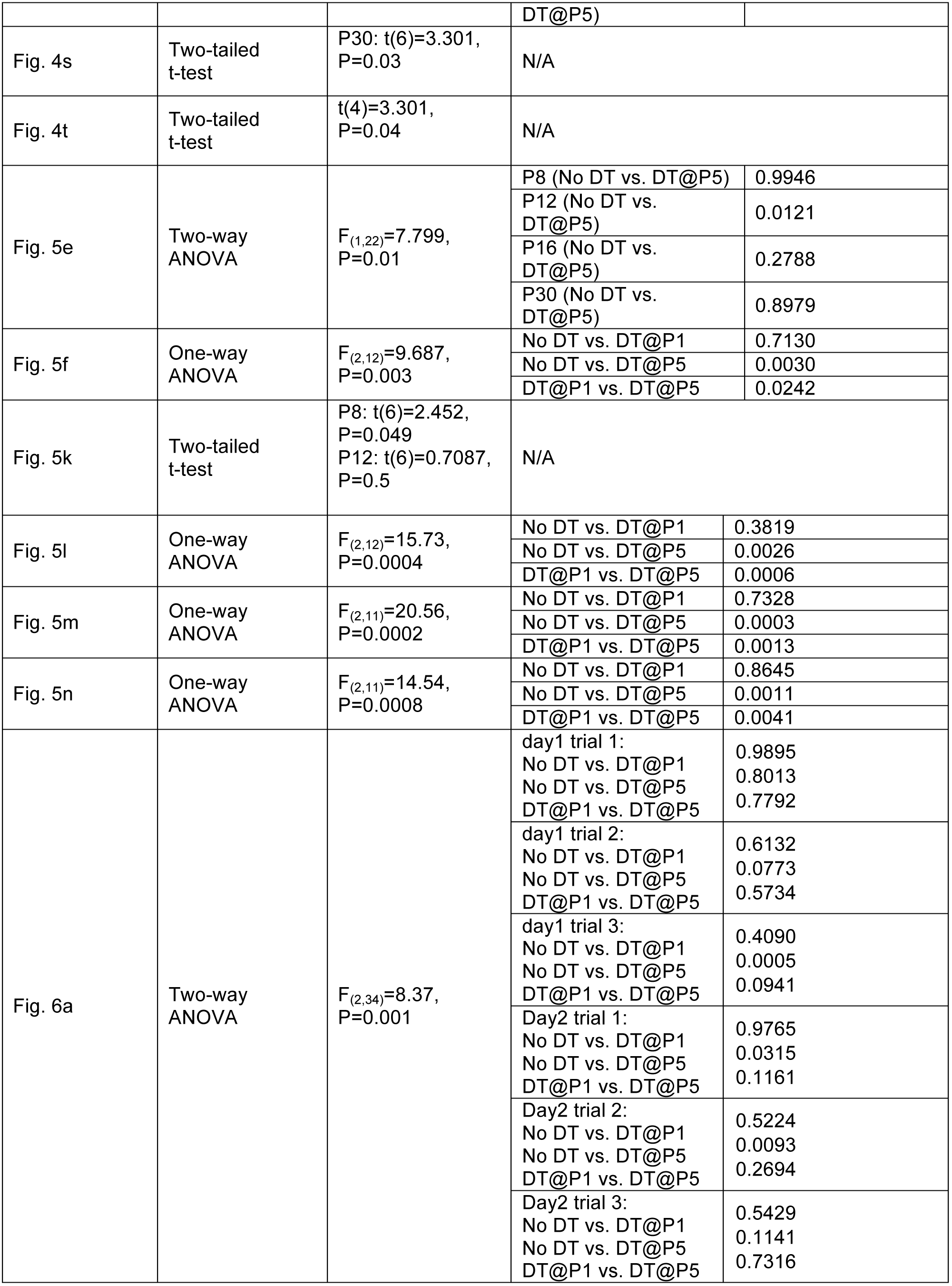
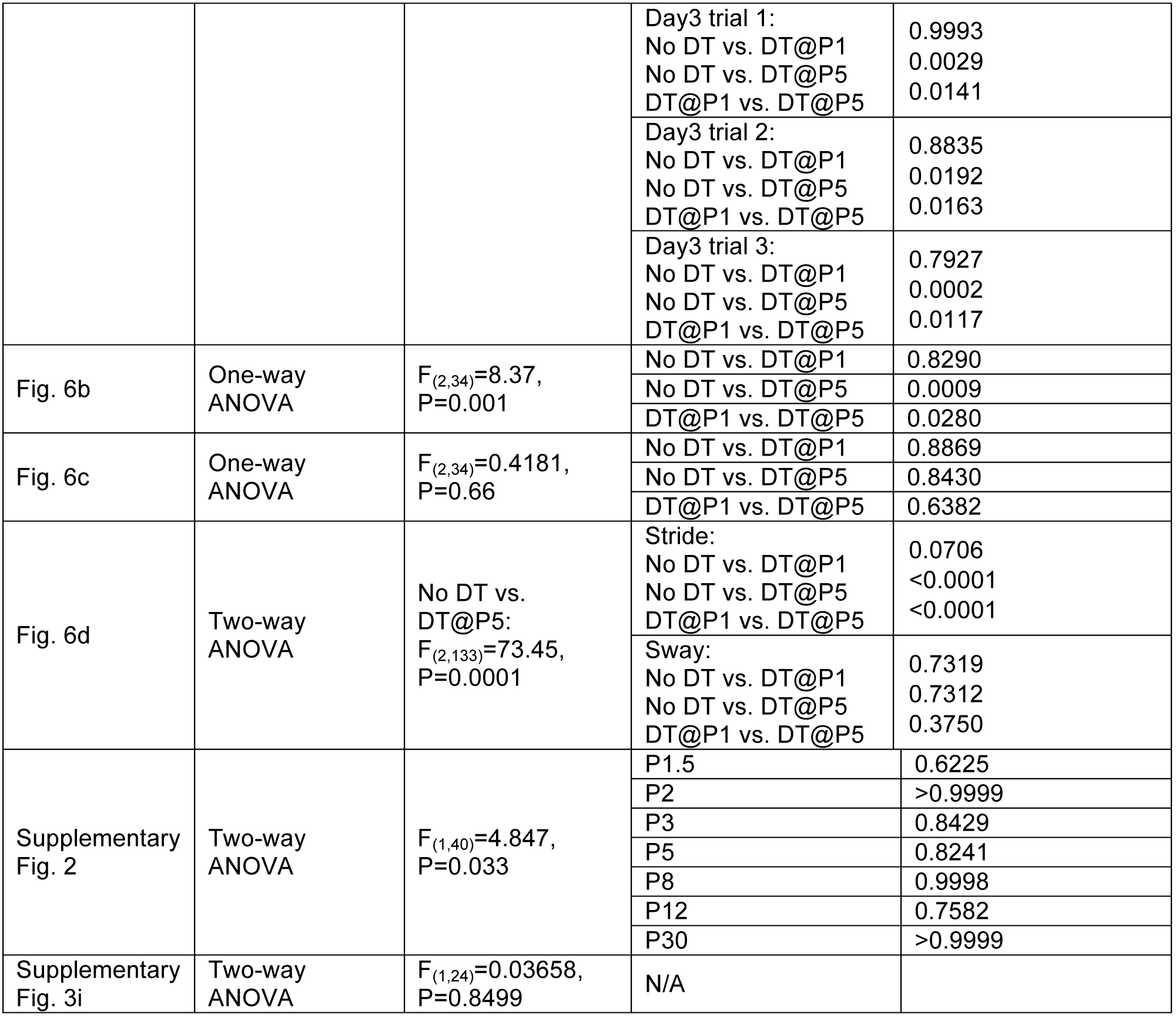
Summary of the statistics performed.

